# Zα and Zβ domains of ADAR1 and ZBP1 bind to G-quadruplexes with low micromolar affinity

**DOI:** 10.1101/2025.11.19.688355

**Authors:** Charles W. Kroft, Jeffrey B. Krall, Michael Warchol, Robb Welty, Alan Herbert, Morkos A. Henen, Beat Vögeli

## Abstract

While it is well established that the Zα domains of ADAR1 and ZBP1 proteins bind Z-form-prone nucleic acids (Z-NAs), it has also been shown that the Zα domain of ADAR1 binds DNA G-Quadruplexes (GQ). However, no binding partner of the structurally homologous Zβ domain of ADAR1 has been identified to date. Based on AlphaFold and molecular dynamics simulations, it has recently been suggested that the Zβ domain of ADAR1 targets its substrate by recognizing GQs. Here, we provide the first experimental evidence for Zβ domain binding to select G-quadruplex RNA and DNA *in vitro*, with structural specificity and low micromolar affinity. We also demonstrate that the Zα domains of ZBP1 bind to both DNA and RNA GQs with similar affinity. These findings extend the range of potential functional roles for these proteins and open new hypotheses for testing in cells.

## 2 Introduction

Z-Binding Domains (ZBDs) are known for their ability to bind Z-conformation nucleic acids (Z-NA) and carry the ability to flip specific right-handed helices of nucleic acids (NA) into Z-NA^1–3^. Detection of double-stranded nucleic acids is an important process occurring in all cells, playing a role in pathways such as innate immune response. Interestingly, both double-stranded DNA and RNA (dsDNA and dsRNA) can adopt higher–energy, left-handed Z-conformation^4–6^, in addition to their common B– and A-conformations, respectively. Z-DNA is generated through supercoiling during transcription and by the ejection of nucleosomes from DNA. In various disease conditions with dysregulated RNA transcription such as viral infection, auto-inflammation, and cancer^7,8^, host retroelements are a significant source of Z-RNA formation^9^. In humans, the only two proteins known to contain ZBDs are adenosine deaminase acting on RNA 1 (ADAR1) and Z-DNA binding protein 1 (ZBP1), which regulate immune responses and/or trigger inflammatory cell death^2,8,10–14^.

ADAR1 is a nucleic acid-binding protein that harbors adenosine-to-inosine editing capability^15,16^. It is expressed in two primary isoforms, the constitutive ADAR1p110 and the interferon-induced ADAR1p150 protein, which contains the Zα domain (Fig. 1A)^17^. While ADAR1p150 undergoes nucleocytoplasmic shuffling, ADAR1p110, is predominantly localized to the nucleus^18^. Both ADAR1p150 and ADAR1p110 incorporate a C-terminal deaminase domain, three intervening dsRNA binding domains (dsRBDs), and the N-terminal enigmatic Zβ domain^16^. Unlike structurally homologous Zα domains, Zβ cannot flip dsDNA and dsRNA into the Z-form or even bind to pre-stabilized Z-NA (Fig. 1D)^19^. Its binding partners and biological functions have remained unknown. Despite this, the Zβ domain is likely functionally important, as its four α-helices are more evolutionarily conserved than the respective Zα domain of ADAR1 across species^20,21^. ADAR1p110 functions to locate and edit self-dsRNA while still in the nucleus, where it is thought to play roles in alternative splicing, miRNA maturation, and recoding of transcripts^22^. The triage of inosine-containing transcripts in the nucleus, mostly of retroelement origin, helps prevent errant immune responses^16,23,24^.

**Figure 1:**
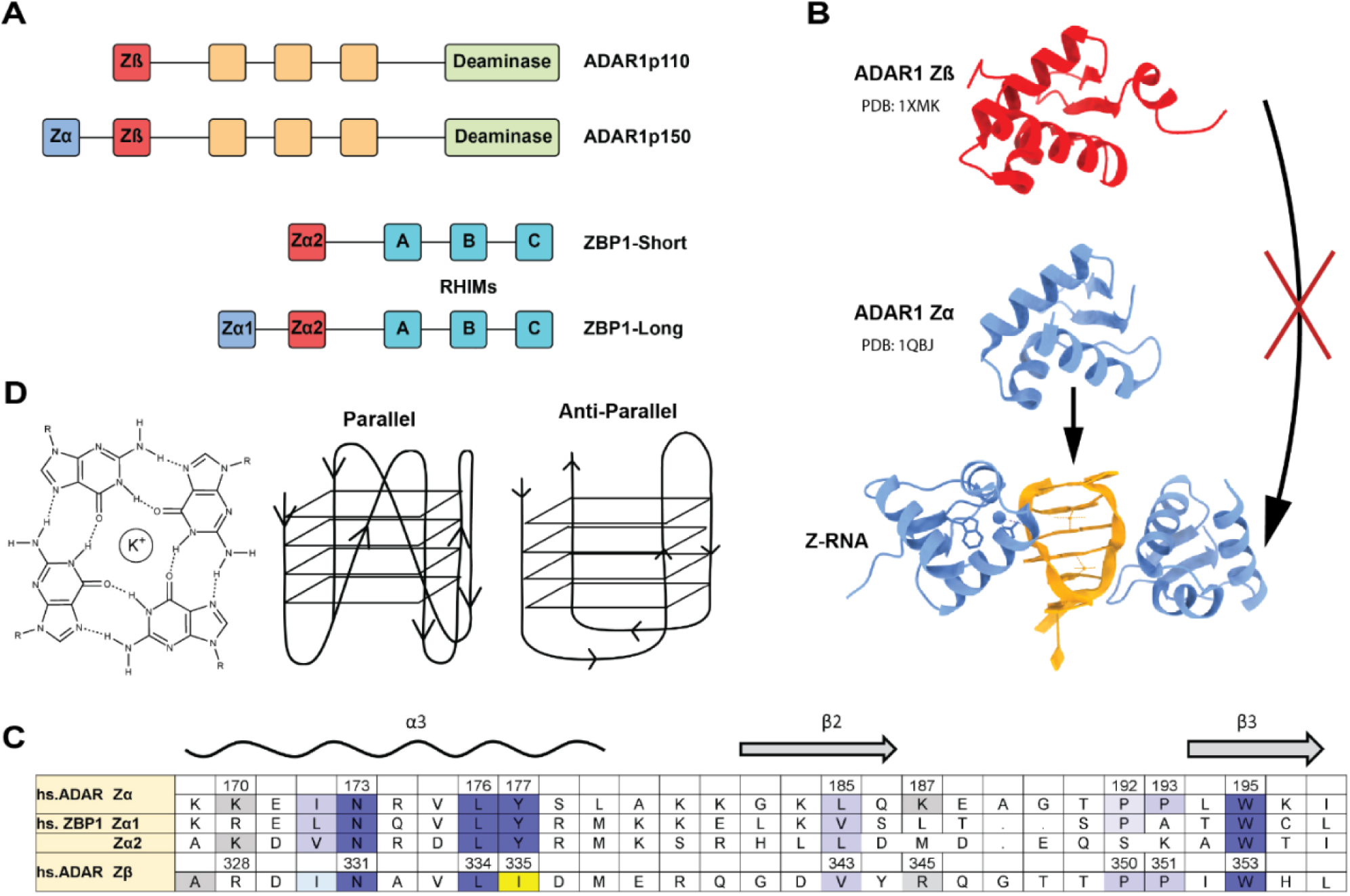
Domain architecture of ADAR and ZBP1 and structure of G-quadruplex. A) Architecture of ADAR and ZBP1 isoforms using the single letter IUPAC coding for amino acids. B) Structure of Zα and Zβ domains, direct interactions with Z-RNA. C) Sequence alignment of human Zα and Zβ domains. Universally conserved Zα family residues are shaded dark blue, with other conservative substitutions shaded light blue. The Y-to-I variant that abrogates Z-NA binding by Zβ is highlighted in yellow. D) G-tetrad, as well as the structure of anti-parallel and parallel GQs.

Both Zα and Zβ are paralogs with distinct evolutionary histories. Both consist of three α-helices and three anti-parallel β-sheets, but Zβ lacks the conserved tyrosine that plays an essential role in ZNA binding^20,25,26^. The Zα NA binding interface is conserved across all members of the Zα domain^2,21,26^. In the Zα domain, N173, Y177, and W195 in the recognition helix (α3) and β-wing are conserved and indispensable (*uniprot*: <u>P55265</u>) for structure-specific interaction with Z-form nucleic acids^25,26^ (Fig 1C).

While ADAR1p110 functions to reduce the level of dsRNA in the cell by promoting inosine-dependent degradation, ADAR1p150 prevents an immune response to endogenous self-dsRNA, most of which are produced by retroelements^27–30^. The recognition of these elements by ADAR1p150 modulates the innate immune response, in part by squelching the activation of inflammatory cell death by ZBP1^11–14^. The full-length product of ZBP1 is comprised of three RIP-homotypic interaction motifs (RHIMs) and two tandem Zα domains (Zα1 and Zα2) (Fig. 1A)^31,32^. The RHIM motifs facilitate downstream signaling upon the interaction of ZBP1 with Z-NA, leading to three major cellular responses: apoptosis, necroptosis, and pyroptosis, depending on various cellular conditions^8^. ZBP1 is spliced into two predominant isoforms, ZBP1-Long (ZBP1-L) and ZBP1-Short (ZBPI-S)^33^. In humans, ZBP1-L contains both Zα domains in addition to the RHIMs^31^. On the other hand, ZBP1-S contains only Zα2 and RHIM motifs. ZBP1-S is known to be sufficient for mediating crisis response, whereas ZBP1-L exhibits a different subcellular localization compared to its shorter counterpart and is more efficient at initiating necroptosis. The findings hint at different roles for the two isoforms, as there are for ADAR1p150 and ADAR1p110^34^. Overall, ZBP1 and ADAR1 function in a regulatory interplay, wherein ZBP1 senses cytosolic Z-NA and dsRNA to initiate innate immune responses and programmed cell death. At the same time, ADAR1 binds and edits these same nucleic acids to suppress aberrant ZBP1 activation and maintain immune homeostasis.

Various targets for Zα domains have been identified in the past, including inverse-repeat ALU (IR-ALU) short interspersed nuclear elements (SINEs) within the noncoding regions of mRNAs^35,36^. Interestingly, it has been shown that Zα also binds to the parallel DNA G-quadruplexes (GQs) formed in the oncogenic c-Myc promoter, albeit with residues distinct from those used for Z-NA binding^37^. Furthermore, it has been demonstrated that ADAR1p110 localizes to telomeres and modulates R-loop formation in an editing-dependent manner^38^. The telomeric repeat-containing RNA (TERRA) produced from telomeric and subtelomeric regions of the genome can activate ZBP1 when a cell is undergoing replicative stress, initiating a MAVS-mediated innate immune signaling and triggering a telomere-driven crisis response^34^. TERRA-colocalized ZBP1 oligomerizes into filaments on the outer mitochondrial membrane as part of the response^34^. However, these interactions were not characterized at the molecular level.

Of interest is that TERRA DNA sequences form anti-parallel GQs in vitro^39,40^. In GQ structures, four guanines basepair to form a G4 tetrad stabilized by Hoogsteen hydrogen bonding. The tetrads then stack on each other to create a four-stranded GQ helix (Fig. 1D)^41–43^. Sequences prone to form GQ are G-rich. The classical GQ motif is composed of four stretches of two or more guanines, with one or more bases serving as loops to connect the four strands (i.e, (G_3_U_x_)_4_). The stability of GQ is also affected by the monovalent ions present in the central cavity. For most GQ folds, K^+^ is the preferred ion, while Li^+^ often disrupts the fold^44,45^. The functions of GQs depend on where they form, but generally, they serve as regulatory and structural elements in key genomic and cellular processes^46,47^.

Recently, we proposed that certain G-rich ALU RNA sequences form GQ structures^48^, serving as a binding target of ADAR1 to localize it to that region of the genome during transcription^49^. As with almost all RNAs, the GQ conformation is parallel and more stable than those formed with DNA^50^. *In silico* molecular docking experiments employing AlphaFold3^51^ and molecular dynamics simulations revealed that a binding interface between both Zα and Zβ to both DNA and RNA GQ structures is plausible, potentially providing an answer to the question of Zβ’s role. In particular, it was proposed that GQs localize ADAR1 to splice sites to offset mis-splicing of pre-mRNA arising from SINE element insertions into active genes, and to contribute to transcript recoding ^52^. Interestingly, GQs are significantly enriched downstream of ALUs, placing them within the RNA loop formed when an ALU inverted repeat folds to create a dsRNA editing substrate. The effects of ADAR1 on splicing are even observed in mice in which the enzyme is catalytically inactive, suggesting that ADAR 1 performs other functions beyond dsRNA editing^53^.

GQs are also enriched in other repeat elements such as LINEs (long interspersed repeats) and the SINE-VNTR-ALUs (SVA) composite hominid retrotransposon, as well as in Human endogenous retroviruses ^52^. Many viruses with GC-rich genomes also harbor GQ-forming sequences^54^. Along with Z-NAs, GQs represent a potential vulnerability that the host can target to modulate the threat they pose.

Given the potential biological importance of the interaction of ADAR1 with GQ to the host, and the previously discovered interaction between Zα and a specific GQ, we seek here to broadly test and characterize the interactions between the ZBDs of both ADAR1 and ZBP1 and various GQ structures.

## 3 Methods

### 3.1 Nucleic acid constructs

The following RNA and DNA constructs were tested for binding to the Zα and Zβ domains: i) TERRA RNA (UUAGGG)_4_ (TERRA-GQ_RNA_), ii) RNA (UUACCG)_4_ (TERRA-mut_RNA_), iii) ALU-RNA GGGA-GGGC-GGGA-GGG (ALU-GQ_RNA_), iv) RNA CCGA-CCGC-CCGA-CCG (ALU-mut_RNA_), v) RNA (GGGU)_3_-GGG (U-loop-GQ_RNA_), and vi) DNA (GGGGTTT)_3_-GGGG (TTT-loop-GQ_DNA_). These sequences are displayed in Table 1.

**Table 1:**
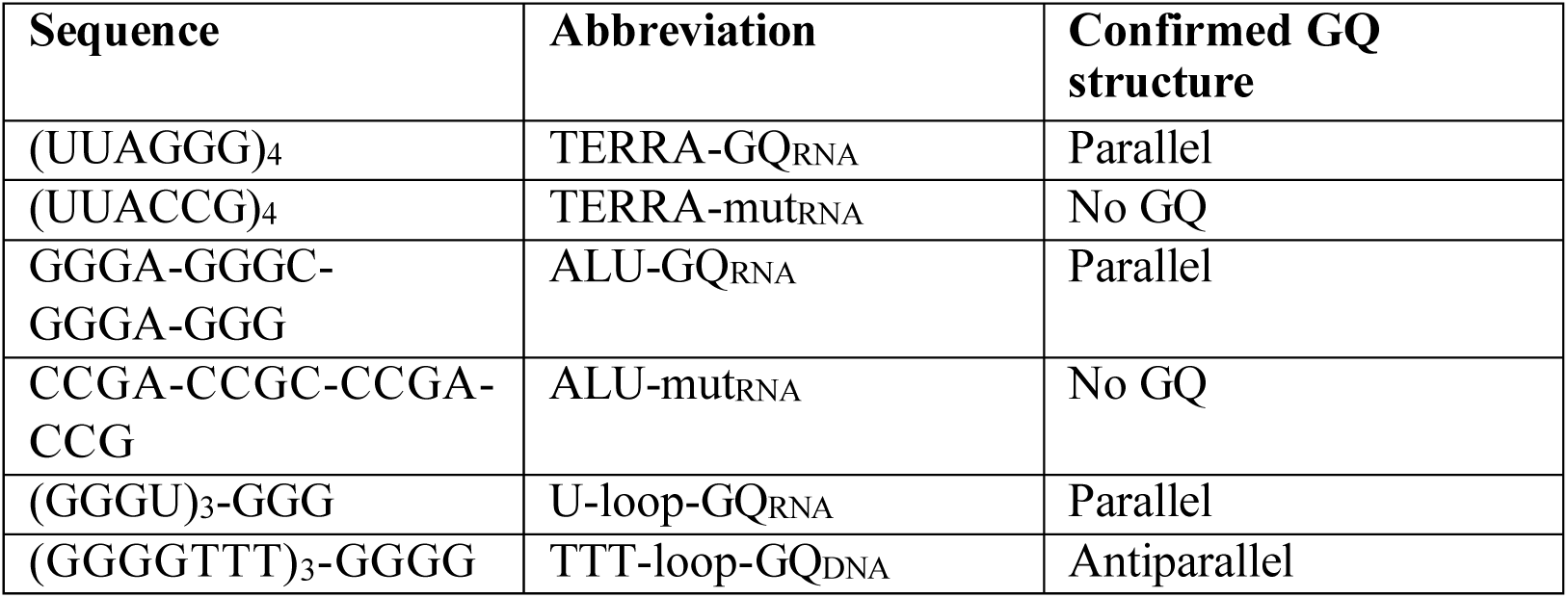
DNA and RNA constructs used in this study.

Each RNA or DNA construct was dissolved in ultra-pure water by diluting to a concentration of 1 mM and heating to 82 °C. Concentrations were confirmed by Nanodrop. To form GQ structures, each RNA or DNA GQ-forming sequence was heat refolded under the following conditions: the NA stock was heated at 82 °C for 10 minutes and allowed to cool down at room temperature for at least 20 minutes. Following this, each construct was diluted to target concentrations in ‘GQ-promoting buffer’ (20 mM NaPi, pH 6.4, 100 mM KCl, 1 mM DTT, 0.5 mM EDTA). Successful GQ folding was confirmed via CD. ALU-mut_RNA_ and TERRA-mut_RNA_ sequences were treated with the same procedure as GQ constructs prior to measurements to ensure experimental consistency.

For GQ ‘unfolding’, the NA constructs were diluted to the target concentration in ‘GQ inhibiting buffer’ (20 mM NaPi, pH 6.4, 100 mM LiCl, 1 mM DTT, 0.5 mM EDTA), heated to 82° C and allowed to cool at room temperature. Additionally, a high NaCl GQ buffer was used, containing 40 mM NaPi, pH 6.4, 100 mM NaCl, 1 mM DTT, and 0.5 mM EDTA.

### 3.2 Protein expression and purification

We derived four protein constructs from the N-terminus of *Homo sapiens* ZBP1 (UniProt: Q9H171): ADAR1 Zα (139-202) and Zβ (291-366), and ZBP1 Zα1 (1–70) and Zα2 (103–166). The codon-optimized constructs were synthesized by Genscript and subcloned into the pET-28a(+) plasmid using NcoI and BamHI restriction sites yielding proteins with an N-terminal 6 × His-tag, a thrombin cleavage site, and a T7 tag. The ordered constructs were expressed and purified as described previously^10,31^. Briefly, the plasmids were transformed and expressed in LOBSTR-BL21(DE3) *E. coli*. The cell cultures for Zα, Zβ, Zα1, and Zα2 were grown in M9 minimal media supplemented with 1 g/L ^15^N ammonium chloride. All cell cultures were grown to an optical density of 0.6 at 600 nm, induced with IPTG at a final concentration of 1 mM, and allowed to express overnight at 21 °C. The cell cultures were centrifuged at 4k RPM for 15 minutes to collect the cell pellets. Pellets were resuspended in the lysis buffer (50 mM Tris–HCl, pH 8.0, 300 mM NaCl, 10 mM imidazole), supplemented with 1 mM dithiothreitol (DTT), and subjected to sonication for cell disruption. The lysate was clarified by centrifugation at 15k RPM for 30 minutes to clear cell debris. The clarified supernatant was applied to 5 × 1 mL HisTrap FF columns (Cytiva, Marlborough, MA), washed with 5 column volumes (CVs) of lysis buffer, 5 CVs of wash buffer (50 mM Tris–HCl (pH 8.0), 1 M NaCl, 10 mM imidazole), and eluted in 1 mL fractions over 3 CVs of elution buffer (50 mM Tris–HCl (pH 8.0), 300 mM NaCl, 500 mM imidazole). The eluents were concentrated to 4 mL, and further purification was completed on a size-exclusion HiLoad 16/600 Superdex 75 pg (Cytiva, Marlborough, MA) in the final GQ NMR buffer (20 mM NaPi, 100 mM KCl, 1 mM DTT, 0. 5 mM EDTA, pH 6.4). ^15^N labeled proteins were concentrated fractions using a 3,000 MWCO concentrator (Millipore-Sigma, Burlington, MA) – Zα to 1.1 mM, Zβ to 2.2 mM, Zα1 to 2.13 mM, and Zα2 to 3.7 mM for later NMR use.

### 3.3 Circular dichroism spectroscopy

For circular dichroism (CD) measurements, the DNA GQ and RNA GQ constructs were prepared at a final concentration of 50 *µ*M in GQ-promoting, inhibiting, or high NaCl buffer. All CD measurements were run using a JASCO J-815 CD spectrometer (using Spectra Manager version 2 (JASCO)) in a 0.1 cm quartz cuvette. For GQ confirmation, ellipticity measurements were taken at 35°C between 320 and 240 nM. Intervals of 1 nM were recorded over three accumulations, and averaged to generate CD spectra, at a rate of 100 nM per minute. To determine the melting temperatures of the GQ sequences, the ellipticity was measured at 264 nM, over a range of 20 to 90 °C.

### 3.4 NMR spectroscopy

For the 1D ^1^H NMR measurements of GQ sequences, the RNA was prepared at a concentration of 100 *µ*M in GQ-promoting buffer as described previously at a volume of 150 µL, including 10 µL of D_2_O to achieve a concentration of 6.6%. The sample was pipetted into a regular non-Shigemi 3 mm NMR tube. For 2D ^15^N-HSQC measurements of ZBDs (ADAR1 Zα and Zβ, ZBP1 Zα1 and Zα2), ^15^N-labeled protein was diluted in GQ NMR buffer to a final concentration of 200 µM at a volume of 150 µL, including 6.6% D_2_O. For binding experiments between nucleic acids and ZBDs, samples were prepared with a final protein concentration of 200 μM and a final RNA concentration of 100 μM, with 6.6% D_2_O. Samples were incubated at 42 °C for 30 minutes before measuring. All RNA and protein NMR experiments were performed at 35 °C.

NMR spectra were acquired on a BRUKER Avance NEO 600 MHz cryoprobe spectrometer equipped with a 5/3 mm triple resonance cryoprobe (CP2.1 TCI) and sample jet, using TopSpin 4.2.0 (Bruker). All ^1^H carrier frequencies were centered on water. 1D NMR experiments were measured using a spectral width of 25 ppm, 256 scans, and 16,384 complex points. The Watergate scheme was used for water suppression.

The ^15^N-HSQC spectra were acquired with a nonuniform sampling (NUS) scheme generated by the NUS@HMS scheme generator^55^ employing 1,024 complex data points in the direct dimension and 50% sampling of the original 200 complex points in the indirect ^15^N dimension. The spectral widths were 13.7 and 35 ppm for the ^1^H and ^15^N dimensions, respectively, with a relaxation delay of 1.3 s and 16 scans. All HSQC spectra were assigned using previously published 3D assignments^10,31^. Data reconstruction of the 2D NUS spectra was performed using the hmsIST software^55^. A solvent subtraction function was applied in the direct dimension. Further data processing and visualization were conducted using NMRPipe/NMRDraw^56^. Spectra analysis was carried out using the CCPNmr analysis software v2.5.1^57^. Except for the titration experiments, all HSQC measurements were recorded with a protein concentration of 200 µM, and a nucleic acid concentration of 100 µM. GQ melting experiments of ALU and U-loop RNA were performed by collecting 1D ^1^H spectra at a temperature range between 35 and 75 °C, with 10 °C intervals. Each experiment was repeated in GQ, promoting, inhibiting, and high NaCl buffers. For NMR titration analysis, a series of ^15^N-HSQC experiments were performed using 200 µM of the protein with a series of nucleic acid concentrations ranging from 0 to 200 µM. The chemical shift perturbation was calculated using the following equation:

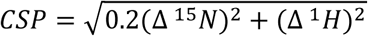

### 3.5 Isothermal titration calorimetry

Nucleic acid and protein constructs were buffer matched by dialyzing twice overnight in the same beaker at 4 °C into GQ-promoting buffer as described above, using 1 kDa cut-off mini dialysis kits (Cytiva). The concentrations after dialysis were measured using a NanoDrop 2000 Spectrophotometer (Thermo Scientific). Protein stocks and nucleic acid stocks were diluted down to a final concentration of 500 µM and 50 μM, respectively. Binding heat was measured on a Malvern ITC200 instrument (run using ITC 200 version 1.26.1 (Malvern)) at 35°C and a stirring speed of 750 rpm, with 180 s injection delays and a reference power of 9 μcal s^-1^. The titration was performed with 19 consecutive 2 μL injections of 500 µM protein into 50 μM nucleic acid, with an initial injection volume of 0.4 μL. All ITC thermograms were fitted using Microcal Analysis version 7 SR4 (OriginLab); the details of the fitting are provided in Freyer and Lewis, 20083. Fitted ITC parameters are shown in Table 3.

**Table 3.** Isothermal titration calorimetry measured parameters.

| Domain | $K_a$ ($\text{M}^{-1}$) | $K_d$ ($\mu\text{M}$) | $\Delta H$ (kcal/mol) | $\Delta S$ (cal/mol/deg) | N (sites) |
| --- | --- | --- | --- | --- | --- |
| ADAR1 $Z\alpha$ | $2.35 \times 10^6 \pm 5.7 \times 10^5$ | $0.43 \pm 0.10$ | $-20.36 \pm 0.72$ | -47.3 | $1.44 \pm 0.03$ |
| ADAR1 $Z\beta$ | $5.54 \times 10^5 \pm 9.2 \times 10^4$ | $1.81 \pm 0.30$ | $-20.05 \pm 1.08$ | -40.2 | $1.12 \pm 0.04$ |

## 4 Results

### 4.1 G-quadruplexes are formed under *in vitro* conditions

We tested the following RNA and DNA constructs for binding to the Zα and Zβ domains (Table 1): i) GQ-forming (UUAGGG)_4_ RNA as found in TERRA sequences (TERRA-GQ_RNA_), ii) a similar RNA sequence (UUACCG)_4_ which is mutated such that it no longer forms a GQ (TERRA-mut_RNA_), iii) GQ-forming GGGA-GGGC-GGGA-GGG RNA as found in ALU elements (ALU-GQ_RNA_), iv) a mutated ALU sequence such that it no longer forms a GQ (CCGA-CCGC-CCGA-CCG) (ALU-mut_RNA_), v) (GGGU)_3_-GGG forming an RNA GQ with minimal loops (U-loop-GQ_RNA_), and vi) (GGGGTTT)_3_-GGGG forming a DNA-GQ (TTT-loop-GQ_DNA_). DNA and RNA constructs are described in Table 1. Prior to conducting binding experiments (see next section), we determined if these nucleic acid constructs form GQ under our *in vitro* conditions.

#### 4.1.1 Circular dichroism confirms the fold of DNA and RNA constructs

TERRA-GQ_RNA_, U-loop-GQ_RNA_, ALU-GQ_RNA_, and TTT-loop-GQ assume parallel (RNA constructs) and anti-parallel (DNA construct) GQ structures in GQ-promoting buffer as expected (Figs. S1A, B). The anticipated spectral characteristics of various nucleic acid structures are listed in Table S1^58,59^. The CD also confirmed that the mutated RNA sequences TERRA-mut_RNA_ and ALU-mut_RNA_ do not adopt a GQ structure, or any alternative structure such as an i-motif (Fig. S1A).

#### 4.1.2 RNA G-quadruplexes are stable to high temperatures

We examined the stability of GQ-forming RNA constructs to further confirm successful folding, as stable GQs are known to have high melting temperature^44,60^. We also confirmed that the predominant buffer ion affects GQ stability in the expected manner^44,45^. In agreement with previous literature, ALU-GQ_RNA_, U-loop-GQ_RNA_, and TERRA-GQ_RNA_ melt at high temperatures in K+ buffer, followed by Na+, with Li+ being the least stabilizing (Fig. S1C). In comparison, we observed that the TERRA-mut_RNA_ construct had a lower melting temperature in both K^+^ and Li^+^ than any other examined RNA construct (Fig. S1C). This indicates successful folding of GQ constructs, and a lack thereof with mutant sequences, enabling us to use ion dependence to refine characterization of the NA interactions with ZBDs and interrogate binding under different buffer conditions.

#### 4.1.3 1D NMR further confirms G-quadruplex structure

Using 1D ^1^H NMR, we further interrogated the fold of GQs by tracking the imino peaks in the 10-12 ppm range, characteristic of a GQ structure^44^. In agreement with the circular dichroism spectra, TERRA-GQ_RNA_ but not TERRA-mut_RNA_ showed imino peaks indicative of GQ structure (Fig. S1D), while ALU-GQ_RNA_ and U-loop-GQ_RNA_ show imino GQ peaks that responded to altering buffer conditions in a manner consistent with the melting experiments (Fig. S1E). NMR visualization of the effect of temperature on GQ structure in various buffer conditions also agreed with previous measurements (Fig. S2).

### 4.2 Zα and Zβ domains interact with G-quadruplexes

To test the binding of the Zα and Zβ domains to the various GQ-forming RNA and DNA constructs, we used 2D Heteronuclear Single-Quantum Coherence spectroscopy (HSQC) NMR, which provides a distinct peak for each ^1^H-^15^N bonded pair in the ZBDs at its respective ^1^H and ^15^N resonances. When the nearby binding of nucleic acids alters the local chemical environment, peaks may shift their frequency positions. In addition, decreasing intensities indicate either a slowed down tumbling caused by increase complex size and/or by exchange (binding and unbinding) on a micro-to millisecond timescale.

#### 4.2.1 Zα and Zβ domains interact with TTT-loop-GQ_DNA_

It has been previously reported that the Zα domain of ADAR1 interacts with a DNA GQ, namely the oncogenic c-Myc promoter parallel GQ^37^. We validated the interaction between Zα and DNA GQs with our anti-parallel TTT-loop-GQ_DNA._ Subsequently, we extended the assays to the Zα1 and Zα2 domains of ZBP1 and the Zβ domains of ADAR1.

Each ZBD-TTT-loop-GQ_DNA_ binding measurement (and all subsequent binding experiments) was conducted at a 1:2 protein-NA ratio. We collected 2D HSQC spectra of both the free and bound proteins to compare binding. Upon addition of the antiparallel GQ DNA to Zα, we observed peak shifts indicative of binding, extending the previous report of an interaction between Zα and a parallel DNA GQ (Fig. 2A).

**Figure 2:**
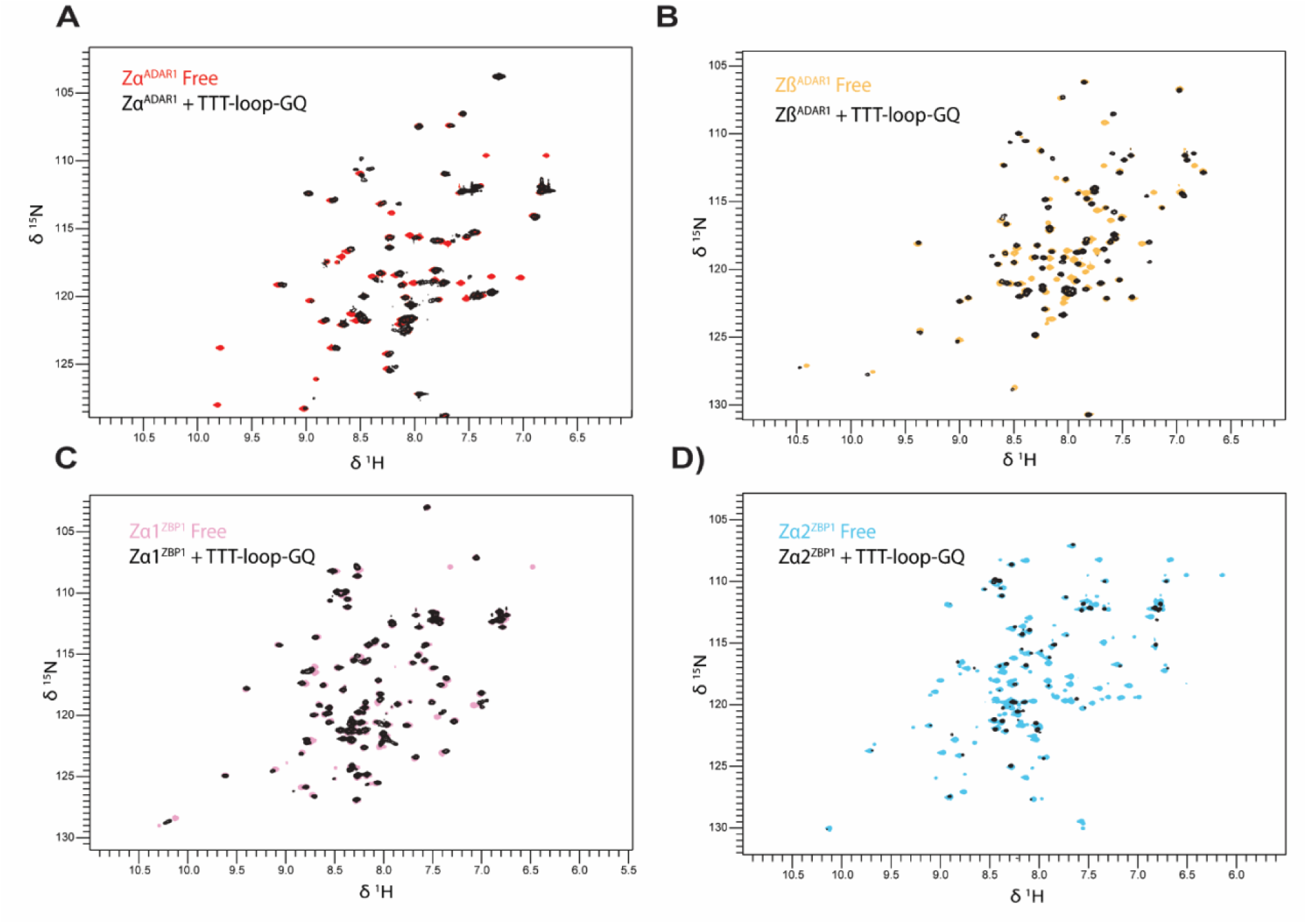
Zα and Zβ domains interact with TTT-loop-GQ_DNA_. Overlaid 2D HSQC spectra of free TTT-loop-GQ_DNA_ and in the presence of A) Zα domain of ADAR, B) Zβ domain of ADAR, C) Zα1 domain of ZBP1, and D) Zα2 domain of ZBP1.

We then examined whether other ZBDs possessed the same capabilities, especially Zβ, whose binding partner was previously unknown. We repeated the binding experiment for Zβ, and for ZBP1 Zα1, and Zα2 domains. All ZBDs interacted with the TTT-loop-GQ_DNA_, showing peak shifts upon addition of the GQ. Zβ had larger shift changes than any Zα domain (see later CSP analyses, sections 4.3.5, 4.4), suggesting potentially tighter binding (Fig. 2B). In contrast, Zα1 appeared to have smaller peak shifts than the other ZBDs (Fig. 2C, Fig. S3E), while Zα2 demonstrated similar peak shifts as did Zβ (Fig. 2D, Fig. S3F).

Peak intensities were also affected by binding, as visualized using peak height ratio plots (bound/free) (Figs. S3A–D). Zα2 displayed the most significant peak height reduction, followed by Zα and Zβ, which were similar, and Zα1 with the smallest reduction. Altogether, these results display binding interactions of all ZBDs with TTT-loop-GQ_DNA_, albeit with potentially different affinities/binding modes as suggested by the varied roles proposed for ZBP1-L and ZBP1-S.

#### 4.2.2 Zα and Zβ domains interact with U-loop-GQ_RNA_

As we established that each of the Zα (ADAR1 and ZBP1) and Zβ domains interacts with the DNA-GQ construct, we next tested if they also interact with RNA GQs. As the simplest RNA structure, we first probed the interactions of ZBDs with the parallel (GGGU)_3_-GGG, whose loops consist of single U nucleotides (U-loop-GQ_RNA_). Upon addition of U-loop-GQ_RNA_ to each ZBD, we observed varying levels of precipitation/phase separation, which decreased upon incubation at 42 °C. Given the homogeneity and reversibility we consistently observed, it is highly likely that, in this case and in all similar instances with other constructs, phase separation prevails. Zα of ADAR1 and Zα2 of ZBP1 experienced the highest level of precipitation, Zα1 of ZBP1 intermediate, and Zβ the least.

When we used HSQC to visualize binding interaction, we saw that upon addition of the U-loop-GQ_RNA_, Zα underwent only slight peak shifts and no visible peak disappearance (Fig. 3A). While this could indicate that Zα does not strongly interact with U-loop-GQ, the observed protein precipitation/phase separation likely means that an interaction is indeed occurring. However, the large protein/RNA complexes so formed prevented HSQC visualization of shifts.

**Figure 3:**
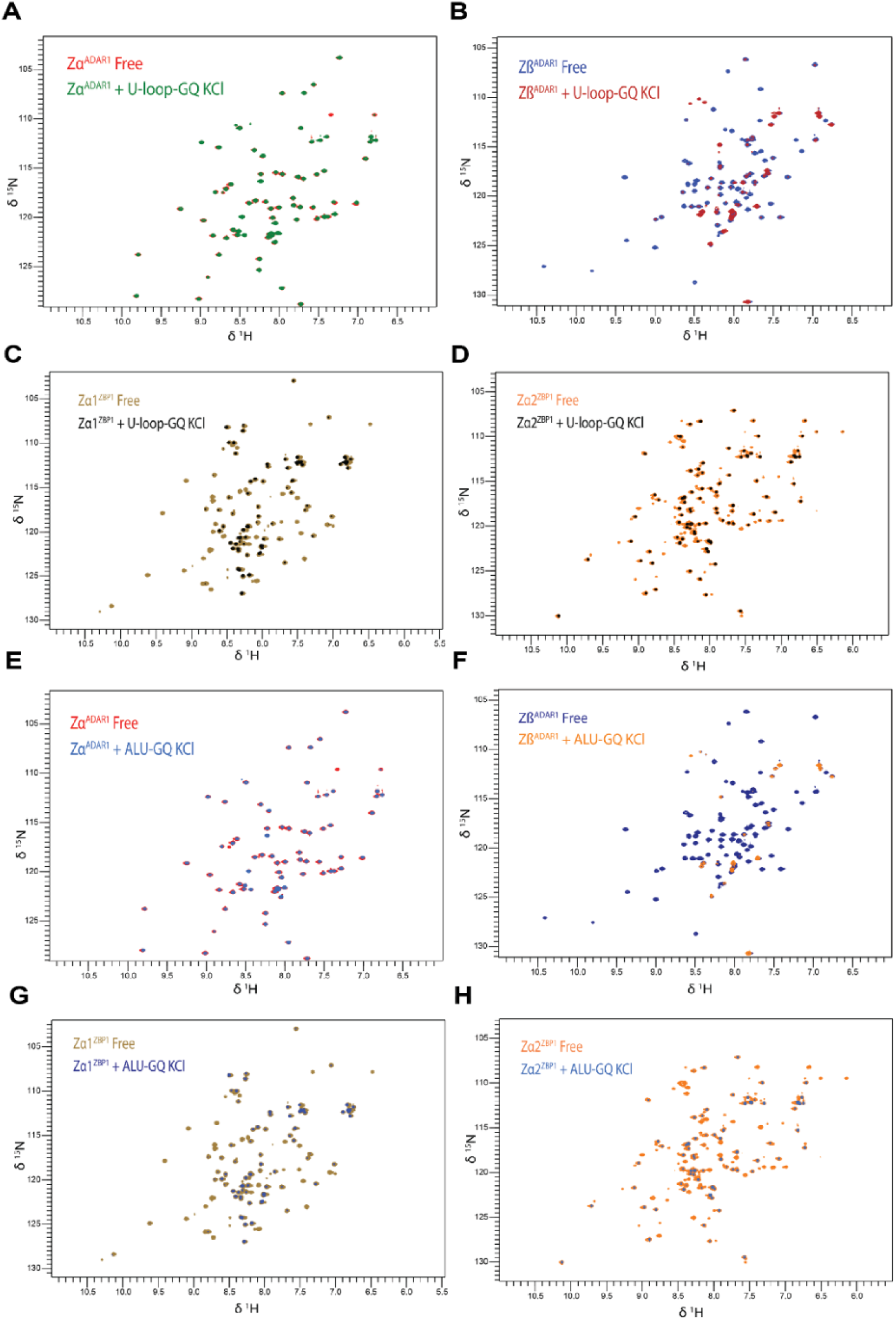
Zα and Zβ domains interact with U-loop-GQ_RNA_ and ALU-GQ_RNA_. Overlaid 2D HSQC spectra of free U-loop-GQ_RNA_ or ALU-GQ_RNA_ and in the presence of A, E) Zα domain of ADAR1, B, F) Zβ domain of ADAR1, C, G) Zα1 domain of ZBP1, and D, H) Zα2 domain of ZBP1.

In contrast, upon addition of the U-loop GQ, Zβ displayed significant peak disappearance, with only minor peak shifting (Fig. 3B). These changes indicate that Zβ does interact extensively with the U-loop-GQ RNA and in a possibly different manner than Zα. Since Zβ precipitated less than Zα did, we can infer that the interaction with U-loop-GQ is different for Zβ than for Zα.

Similarly, Zα1 showed significant peak disappearance and a reduction in peak height, indicative of binding to U-loop-GQ. Only minor peak shifts were observed (Fig. 3C). These results are similar to the spectrum for Zβ. Intriguingly, both Zβ and Zα1 showed less precipitation than Zα or Zα2.

For Zα2, we observed similar levels of protein precipitation as Zα. As expected, we observed an HSQC spectrum with only minor peak losses and minimal shifts (Fig. 3D). This result is likely again due to complex formation, which obscured the visualization of shifts.

We expected to see a direct correlation between the peak height reductions and degree of precipitation; intriguingly, though, we saw the opposite trend, with Zα and Zα2 showing the least peak height reduction relative to free protein, followed by Zα1, and finally Zβ with the greatest degree of peak reduction (Fig. S4). This could be explained by phase separation, which sequesters nucleic acids, leaving mainly unbound protein in solution. In the absence of phase separation, a higher fraction of protein and nucleic acid remain in a free state, allowing visualization of their interactions. Thus, peak height reductions appear to reflect the fraction of nucleic acid interacting with protein and the strength of binding, rather than arising from the formation of condensates.

Overall, these HSQC spectra indicate clear binding interactions between the Zβ and Zα1 domains with the U-loop-GQ_RNA_, as both spectra displayed extensive peak disappearances. While we did not observe peak shifts for the ADAR1 Zα and ZBP1 Zα2 domains, we did observe significant reductions in peak intensity for both, but less pronounced than those for ADAR1 Zβ and ZBP1 Zα1.

#### 4.2.3 Zα and Zβ domains interact with ALU-GQ_RNA_

Given that each ZBD binds to a canonical RNA parallel GQ, we next tested the interactions with a more biologically relevant construct, ALU-GQ_RNA_. As ADAR1 is known to localize to regions of the genome replete with ALU SINE repeats, it provided an appealing motif to study. Notably, our recent study proposed that both Zα and Zβ could bind to a GQ of a specific RNA element found in ALU^49^. We designed an ALU-GQ_RNA_ construct with this element in mind, as it is potentially a highly prevalent, biologically relevant ADAR1 target.

With the ALU-GQ_RNA_, complex precipitation was more pronounced than with the U-loop-GQ, with all four ZBDs affected equally. ADAR1 Zα showed no apparent peak shifts or disappearance, but only peak height reductions (Fig. 3E, Fig. S 5A). The peak height reductions were more intense than those observed for the U-loop-GQ_RNA_, and less than those of other ZBDs (Fig. S5).

Zβ, unlike Zα, showed extensive peak disappearance with ALU-GQ_RNA_ (Fig. 3E). The peak disappearance was more pronounced than that seen with U-loop-GQ and is the greatest of any seen for the ZBD interactions with ALU-GQ_RNA_ (Fig. S 5B). We observed greater peak disappearance in HSQC and greater peak-height reduction. This result is highly significant, given that Zβ precipitation levels were lower than or equal to those of Zα.

Zα1 displayed very similar spectra with ALU-GQ_RNA_ and U-loop-GQ_RNA,_ showing numerous peak disappearances (Fig. 3F). Curiously, despite seeing varying degrees of precipitation between the two constructs, the spectra remain nearly identical, and the average peak height reduction between these two experiments was nearly equivalent (Fig. S 5C). These observations suggest that Zα1 binds similarly to both parallel strand GQ RNAs.

Zα2 demonstrated a different interaction with ALU-GQ_RNA_ than with U-loop-GQ_RNA_; despite similar levels of precipitation, we observed significant peak height reductions with ALU-GQ_RNA_ that were not seen with U-loop-GQ_RNA_ (Fig. 3H, Fig. S 5D).

The results reveal that the interactions of the four ZBDs with the ALU-GQ_RNA_ are similar to those with U-loop-GQ_RNA_, but not equivalent. We also observed greater precipitation with Zβ and ALU-GQ_RNA_ than with U-loop-GQ_RNA_. Despite this outcome, both GQs produced similar large and specific changes to the Zβ spectra and peak height. Zα2 also showed a significant difference between the two GQs, with greater peak disappearance with ALU-GQ_RNA_ than with U-loop-GQ_RNA_ despite a similar level of precipitation. In contras t, Zα and Zα1 showed similar spectra for both U-loop-GQ_RNA_ and ALU-GQ_RNA_. Overall, for both RNA GQs, Zβ underwent the most significant changes, followed by Zα2, Zα1, and lastly Zα.

#### 4.2.4 Zα and Zβ domains interact with TERRA-GQ_RNA_

We also examined the binding interaction between ADAR1 subunits and the parallel-stranded TERRA-GQ_RNA_ construct. As all ZBDs can interact with GQs, we focused on ADAR1, motivated by the previously published hypothesis proposing specific Zβ interactions with GQs. The TERRA-GQ_RNA_ construct is taken from the telomeric repeat-containing RNA and forms a G-quadruplex under our experimental conditions (Fig. S1A). Localization of both ADAR1 and ZBP1 to telomeric regions has been reported previously^34,38^. However, the interactions have not been previously characterized at the molecular level. As a well-established G-quadruplex, the TERRA-GQ_RNA_ also serves as a positive control for our other constructs.

While showing extensive precipitation, Zα underwent very extensive peak disappearance, indicative of tight binding to the TERRA-GQ_RNA_ (Fig. S6). These effects are in stark contrast to those previously seen for Zα with other GQ: the TTT-loop-GQ_RNA_ resulted primarily in shifts, while U-loop-GQ_RNA_ and ALU-GQ_RNA_ caused only peak height reductions. Thus, TERRA-GQ_RNA_ presents a unique binding interaction for Zα. Peak heights were almost entirely extinguished, rivaling reductions seen for Zβ with previous RNA GQs (Fig. S7A). It may be worth noting that the two GQs resulting in visible HSQC modifications for Zα (TTT-loop-GQ_DNA_ and TERRA-GQ_RNA_) are 25 and 24 nucleotides in length, whereas the shorter constructs (ALU-GQ_RNA_ and U-loop-GQ_RNA_ are 15 nucleotides each) show no changes to the spectra. This could be due to the inherently lower stability of the longer constructs, or a preference for Zα to bind longer nucleic acids. In contrast, binding of Zα to ZNA requires a binding site of only 6 bps^61^.

Zβ also showed significant peak disappearance and shifts upon TERRA-GQ_RNA_ addition (Fig. S6B). Similar to Zα, we observed considerable precipitation. The chemical shift perturbations were large, and paired with large peak disappearance, indicating extensive binding (Figs. S7C, E). These results corroborate previous findings, as both domains readily bind to GQs, but are unique in their interactions with different GQ constructs.

### 4.3 ZBPs interact with a broad range of nucleic acid folds

Zα domains are known to bind to Z-NA, as well as A-RNA and B-DNA, albeit with weaker affinity^62^. In contrast, Zβ domains have not been shown to bind any nucleic acids to date. While we have shown that all ADAR1 and ZBP1 ZBDs bind to GQs, their interactions appear to depend on GQ architecture. Therefore, we sought to investigate whether these domains bind nucleic acid motifs other than GQs (and A-, B-, and Z-form double-stranded nucleic acids in the case of Zα domains), to determine whether ZBDs do actually bind GQs with greater specificity.

#### 4.3.1 ADAR1 and ZBP1 ZBDs interact with disrupted G-quadruplexes

To assess if Z-domains bind GQs with structure-specific recognition, we generated mutant ALU and TERRA constructs that interrupt the formation of G-tetrads and therefore prevent G-quadruplex formation (Fig. S1A). Mutation of each GGG repeat to CCG (TERRA-mut_RNA,_ ALU-mut_RNA_) successfully disrupted the GQ structure (Fig. S1A), yielding RNA expected to be predominantly single-stranded.

Zα exhibited extensive peak shifts with both mutant constructs, in contrast to the previous results, which showed only peak disappearance. TERRA-mut_RNA_ induced larger shifts than ALU-mut_RNA_ (Figs. 4A, E, S 7 E, S 9A). Zα still exhibited phase separation/crashing out with the ALU-mut_RNA_, albeit to a lesser extent than seen with U-loop-GQ_RNA_ or ALU-GQ_RNA_. This may indicate that binding was occurring over a different time regime or with different dynamics. Notably, we observed greater peak-height reduction here than with any previous construct, except TTT-loop-GQ_DNA_ (Fig. S8A), which is counterintuitive given that precipitation was less pronounced. TERRA-mut_RNA_ binding yields even greater levels of peak height reduction than with ALU-mut_RNA_, second only to TERRA-GQ_RNA_.

**Figure 4:**
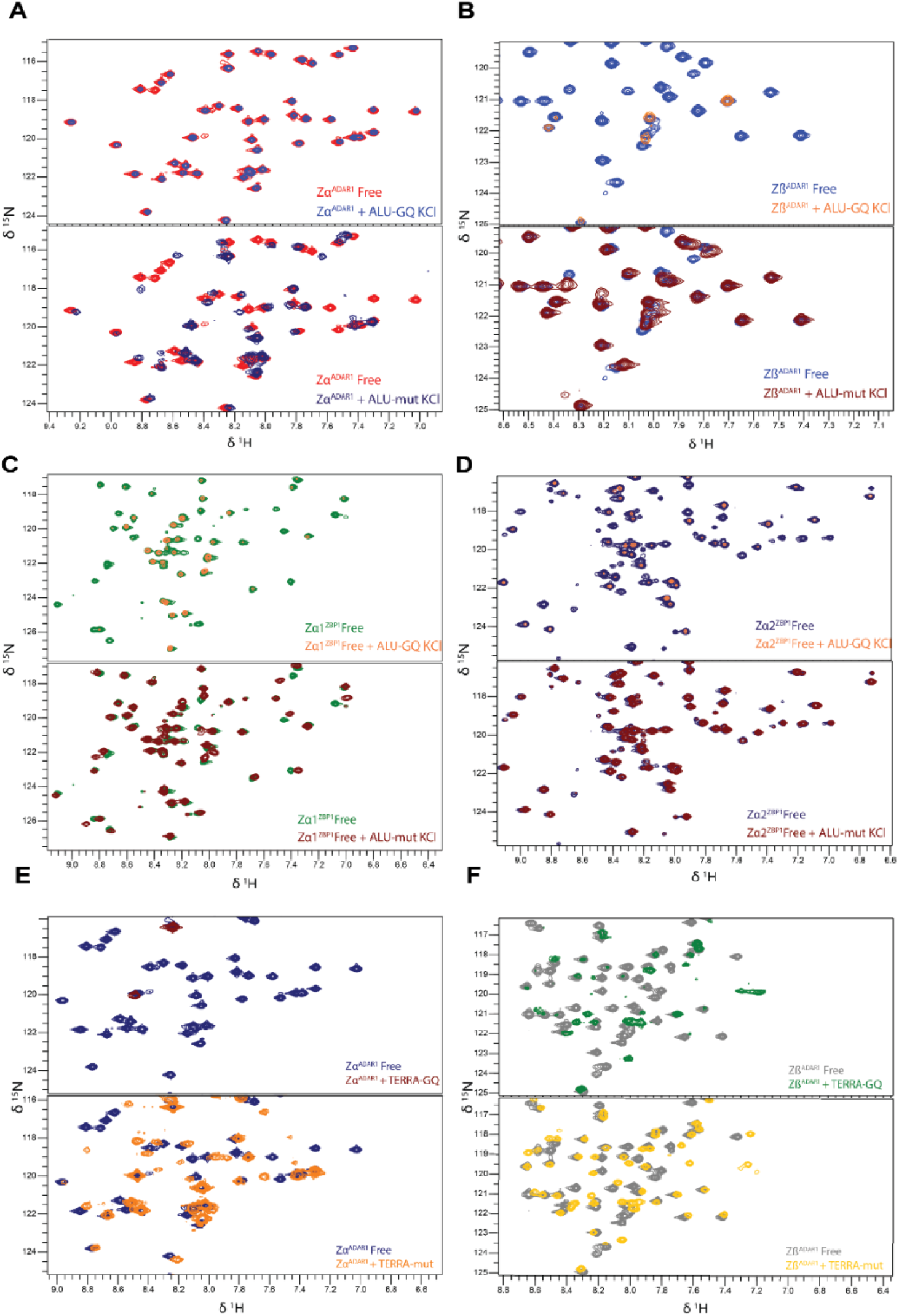
Comparison of Z-domain binding to GQs or disrupted counterparts. Overlaid 2D HSQC spectra of free ALU-GQ_RNA_ or ALU-mut_RNA_ and in the presence of A) Zα domain of ADAR1, B) Zβ domain of ADAR1, C) Zα1 domain of ZBP1, and D) Zα2 domain of ZBP1. Overlaid spectra of TERRA-GQ_RNA_ or TERRA-mut_RNA_ in the presence of E) Zα of ADAR1 and F) Zβ of ADAR1.

Zβ also bound both constructs, which primarily resulted in peak shifts (Figs. 4B, 4F, S7F, S9B). Intriguingly, titrating ALU-mut_RNA_ to Zβ resulted in no precipitation, but we observed less peak disappearance compared to our other GQs (Figs. 4B, S8B). TERRA-mut_RNA_ causes larger shifts than those seen for ALU-mut_RNA_. However, TERRA-GQ_RNA_ caused a markedly greater reduction of peak intensity than TERRA-mut_RNA_ (Figs. S7C, D). Taken together, Zα and Zβ bind to both G-quadruplex and single-stranded RNA constructs, with both ZBDs binding with different modes/dynamics to these various structures.

Zα1 behaved similarly to Zβ when bound to ALU-mut_RNA_; minimal precipitation was observed. While ALU-GQ_RNA_ induced significant peak disappearance with Zα1, ALU-mut_RNA_ primarily yielded peak shifts, and caused less peak height reduction than any G-quadruplex tested with Zα1 (Figs. 4C, S9C).

Zα2 displayed a distinct interaction with ALU-mut_RNA_. We observed substantial phase separation upon addition of the ALU-mutRNA, and the resulting HSQC spectra showed marked reductions in peak intensity with minimal chemical shift changes (Figs. 4D, S8D, S9D).

These results indicate that ZBDs are capable of binding nucleic acids beyond canonical duplex DNA/RNA, Z-form motifs, and GQs. However, the interactions with ALU and TERRA mutant sequences differ from those with their GQ counterparts. In some cases (e.g. Zβ and Zα1), the interactions appear weaker than those observed with wild-type GQs, whereas for Zα, the spectral changes are notably more distinct than those seen with the GQs.

#### 4.3.2 Buffer ion affects ADAR1 Zβ but not Zα binding to ALU and U-loop-GQ

While our previous measurements were conducted in GQ promoting K^+^ buffer, the following experiments were performed in Li^+^ buffer to assess whether interactions reflect structural or sequence specificity. As shown, the buffer ion markedly influences the stability of GQ, with K^+^ having the strongest stabilizing effect, and Li^+^ the weakest (Figs. S 1C, S1E, S2). W e hypothesized that modulating the predominant ion in buffer would allow further examination of the specificity of ZBD-G-quadruplex interactions.

As Zα exhibited no significant changes to its spectrum when interacting with ALU-GQ_RNA_, it is unsurprising the spectra were similar with KCl and LiCl buffers. There were no significant differences in peak intensity between these two, either (Figs. S5A, S11B). These observations suggest that Zα does not exhibit a strong preference for the stability or the fold state of ALU-GQ_RNA_. We saw similar interactions with U-loop-GQ_RNA_, where neither buffer condition induced large peak disappearance or shifts (Figs. S10 B). In fact, we see slight decreases in our peak heights in the Li^+^ over the K^+^ sample (Figs. S4A, S11A), although these variations were minor.

Intriguingly, Zβ exhibited a strong response to buffer ion. While we saw extensive peak disappearance with ALU-GQ_RNA_ and U-loop-GQ_RNA_ in KCl, the vast majority remained visible in Li^+^ (Figs. S10C, D). These changes, together with a large relative increase in peak height (Figs. S4B, S5B, S11C, D), suggest a strong difference in binding between buffer conditions. U-loop-GQ_RNA_ exhibited the same effects, but to a slightly lesser extent. Overall, these data indicate that Zβ is sensitive to buffer ion when interacting with GQs, implying that G-quadruplex stability—and potentially its structure—modulates binding.

#### 4.3.3 ADAR1 Zα and Zβ show low micromolar binding affinities to G-quadruplexes

Given the trends observed in the NMR spectra of the ZBDs upon addition of various nucleic acid constructs, the typical Kd appears to fall in the low micromolar range, possibly even the high nanomolar range. However, because peak disappearance is often the primary effect of binding, a more refined quantification by NMR titration experiments is challenging for many interactions studied here. Therefore, we conducted select isothermal titration calorimetry (ITC) experiments and, when feasible, extracted microscopic Kd values using NMR titration experiments.

We were able to extract affinities for both ADAR1 ZBDs with TERRA-mut_RNA_. These are microscopic affinities that provide residue-by-residue values. *K*_d_ values were determined by fitting peak position changes, that is, chemical shift perturbation (CSP) as a function of ligand concentration. We obtain overall affinities of approximately 108 µM for Zα and 42 µM for Zβ (Figs. 5A, B).

**Figure 5:**
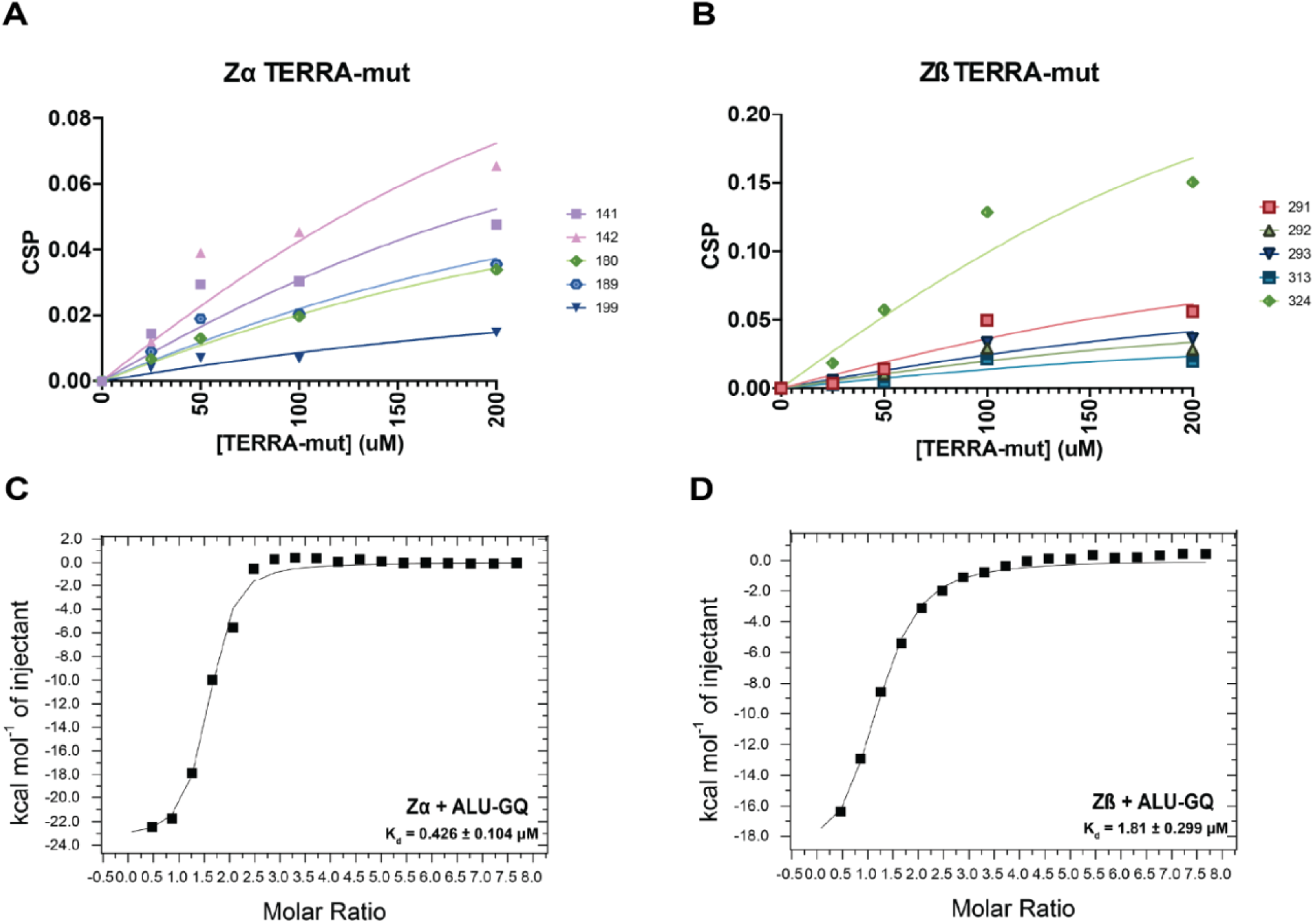
Affinity analysis of interaction of ADAR1 Zα and Zβ subunits with TERRA-GQ_RNA_ and TERRA-mut_RNA_. Residue-specific CSP-based binding affinity curves for A) Zα and B) Zβ binding to TERRA-mut_RNA_. Highlighted residues shown in legend. ITC curves of C) Zα domain and D) Zβ domain binding to ALU-GQ_RNA_.

Interactions that primarily result in peak disappearance are likely to reflect stronger affinities. To determine these values, we used ITC to measure the affinity between both ADAR1 ZBDs with ALU-GQ_RNA_, given the potential role this fold plays in targeting ADAR to ALU repeat editing substrates. Unexpectedly, we saw low-micromolar affinity for both Zα and Zβ when binding to this G-quadruplex. Zα measured a *K*_d_ of 0.43 ± 0.10 µM, with an enthalpy of –23.56 ± 0.72 kcal/mol and a calculated entropy value of –47.3 cal/mol/deg. Zβ measured a *K*_d_ value of 1.81 ± 0.30 µM, enthalpy of –20.48 ± 1.08 kcal/mol, and enthalpy value of –40.2 cal/mol/deg (Figs. 5C, D). These affinities are approximately one order of magnitude larger than their disrupted counterparts. The results suggest that binding of ADAR Zα and Zβ to parallel GQ RNAs GQs is both specific and of high affinity. These values are summarized in Table 3.

As noted, there are ligand-specific variations in the binding of different ZBDs to various GQ folds. Taking the peak disappearance/peak shifting patterns as a proxy for the affinities (see also Table 4), the data imply that all RNA GQs may have a *K*_d_ in the low-micromolar range. In contrast, the affinity for anti-parallel-strand DNA G-quadruplex is one order of magnitude weaker, similar to that of disrupted RNA GQs. It is important to note that these represent macroscopic affinities. The values measured by ITC may reflect a summation of binding events, including specific substrate-ligand interactions and non-specific, multivalent interactions known to occur within phase separates.

**Table 4.** Type of interaction observed in HSQC NMR spectra.

| Domain | U-loop-GQ<sub>RNA</sub> | ALU-GQ<sub>RNA</sub> | ALU-mutRNA | TERRA-GQ<sub>RNA</sub> | TERRA-mutRNA | TTT-loop-GQ<sub>DNA</sub> |
| --- | --- | --- | --- | --- | --- | --- |
| ADAR1 Zα | PI | PI | CS | PI | CS | CS |
| ADAR1 Zβ | PI | PI | CS | CS, PI | CS | CS |
| ZBP1 Zα1 | PI | PI | CS | - | - | CS |
| ZBP1 Zα2 | PI | PI | PI | - | - | CS |

CS = chemical shift changes; PI = peak intensity changes

#### 4.3.4 Zβ exhibits a preference for TTT-loop-GQ_DNA_ while Zα does not

While both Zα and Zβ showed significantly better binding to an RNA GQ than ssRNA, this is not the case for all constructs, notably TTT-loop-GQ_DNA_. Despite extensive peak disappearance in the HSQC spectra of the ADAR1 ZBDs in complex with many constructs (Zα with TERRA-GQ_RNA_, Zβ with ALU-GQ_RNA_, U-loop-GQ_RNA_, TERRA-GQ_RNA_), other constructs instead show CSPs with varying degrees of peak shifts. For Zα, we obtained measurable CSPs for both mutant sequences and the TTT-loop-GQ_DNA_, as well as minor but consistent shifts for U-loop-GQ_RNA_ (which likely binds tightly, as indicated by the substantial peak height loss). Notably, certain residues consistently shifted in the same direction upon the addition of nucleic acid (Fig. 6A). This consistent directionality and ordering of shifts suggest that these residues are specifically interacting with nucleic acids, providing a means to investigate Zα’s preference across constructs. Comparing these shifts to titration curves with TERRA-mut_RNA_, which show similar directional shifts as a function of ligand concentration, supports the hypothesis that for each construct, peak shift magnitude is likely correlated with the binding strength (Fig. S12).

**Figure 6:**
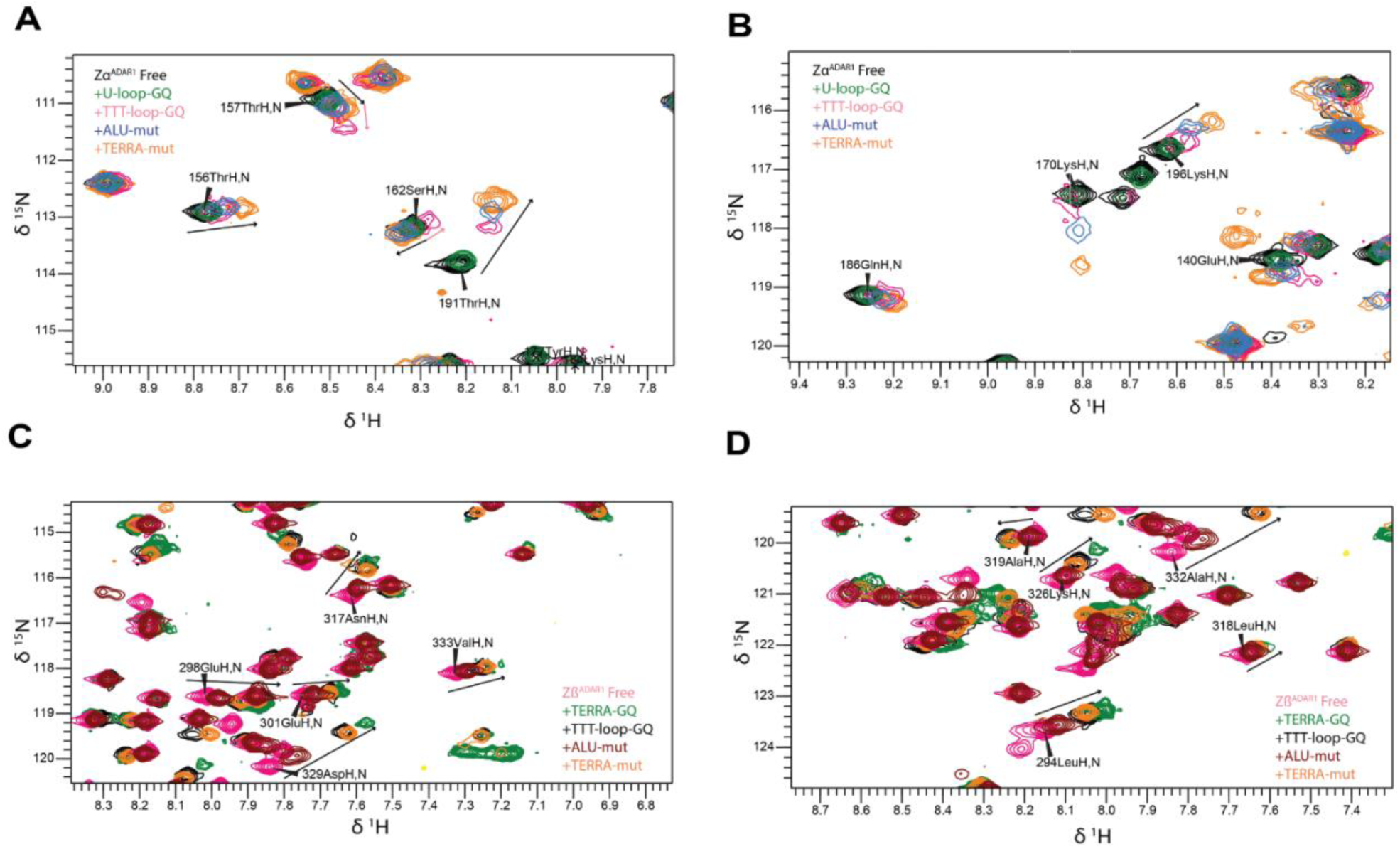
DNA and RNA constructs induce varying peak shift changes in ADAR1 Zα and Zβ. A, B) Extended regions of overlaid HSQC spectra of free Zα domain and in complex with TERRA-mut_RNA_, ALU-mut_RNA_, TTT-loop-G_DNA_, and U-loop-G_RNA_ are shown. C, D) Extended regions of overlaid HSQC spectra of free Zα domain and in complex with TERRA-mut_RNA_, ALU-mut_RNA_, TTT-loop-G_DNA_, and U-loop-G_RNA_ are shown.

When comparing constructs with Zα, a trend emerges: the two non-GQ constructs (TERRA-mut_RNA_ and ALU-mut_RNA_) consistently exhibit the greatest magnitude of shifts, whereas the two GQ constructs with visible CSPs (TTT-loop-GQ_DNA_ and U-loop-GQ_RNA_) exhibit smaller shift changes (Figs. 6A, B). This suggests that Zα preferentially binds non-GQ nucleic acids over these specific GQs, particularly TTT-loop-GQ_RNA_. However, certain residues deviate from this trend: some display shifts for TTT-loop-GQ_DNA_ but not for the other three constructs or shift in a different direction than observed for the other sequences (residues 157, 162) (Fig. 6B). This may reflect residues in Zα that interact differently with GQ versus non-GQ nucleic acids, mediating specific DNA interactions, or due to the fact that TTT-loop-GQ_DNA_ is the only anti-parallel GQ in this study.

We also examined peak shifts for Zβ, as binding to some GQ constructs retained sufficient peak intensity to analyze CSPs. We compared TERRA-mut_RNA_, ALU-mut_RNA_, TTT-loop-GQ_DNA_ and TERRA-GQ_RNA_. A consistent trend emerged here: TERRA-GQ_RNA_ peaks always shifted further than or equal to the other constructs; TTT-loop-GQ_DNA_ shifts were next, consistently greater than or equal to those of TERRA-mut_RNA_ and ALU-mut_RNA_. TERRA-mut_RNA_ exhibited the next greatest magnitude of shifts, with ALU-mut_RNA_ showing the least (Figs. 6C, D). This trend contrasts Zα, where Zβ exhibited an apparent preference for these specific GQs compared to our non-GQ nucleic acids. Taken together, these results indicate that Zβ has a preference for DNA/antiparallel GQs, while Zα has a preferential binding to ssRNA over specific GQ constructs.

### 4.4 ADAR1 domains bind to G-quadruplexes primarily via α-helices

Having observed that specific residues in both Zα and Zβ exhibit consistent CSPs upon interaction with GQs, we can extract information regarding the binding interface with GQ nucleic acids. In principle, similar information could be obtained from peak height intensity plots. However, for constructs that show no peak shifts and only peak disappearance, it is not possible to distinguish residues involved in direct interaction from those peaks that disappear due to slowed overall tumbling. Therefore, we analyzed only spectra showing CSPs to investigate the binding interface. Significant chemical shifts were defined as (*ShiftDist − Mean*)*/STD >* 1.

We compared our Zα binding site results with those presented in a previous study that demonstrated interactions with a DNA G-quadruplex^37^. In that study, Zα engaged the G-quadruplex primarily through α-helices 2 and 3 and the β-wing, affecting residues Ala158, His159, Lys169, Lys170, Glu171, Asp173, Arg174, Val175, Lys176, Lys182, Ala189, and Thr191. These residues substantially overlap with those that mediate Z-NA binding, for which Tyr177 and Trp195 are also critical. Based on the CSPs of the TTT-loop-GQ_DNA_ and U-loop-GQ_RNA_ (Fig. 7A, B), we found that the primary residues interacting with our GQs are Glu140, Lys170, Glu171, Ile172, Asp173, Arg174, Val175, Leu176, Tyr177, Ser178, and Thr191. This set is highly similar to that reported previously for G-quadruplex binding, and also overlaps extensively with the Z-NA interaction interface. These residues similarly map to α-helix 3 and the β-wing.

**Figure 7:**
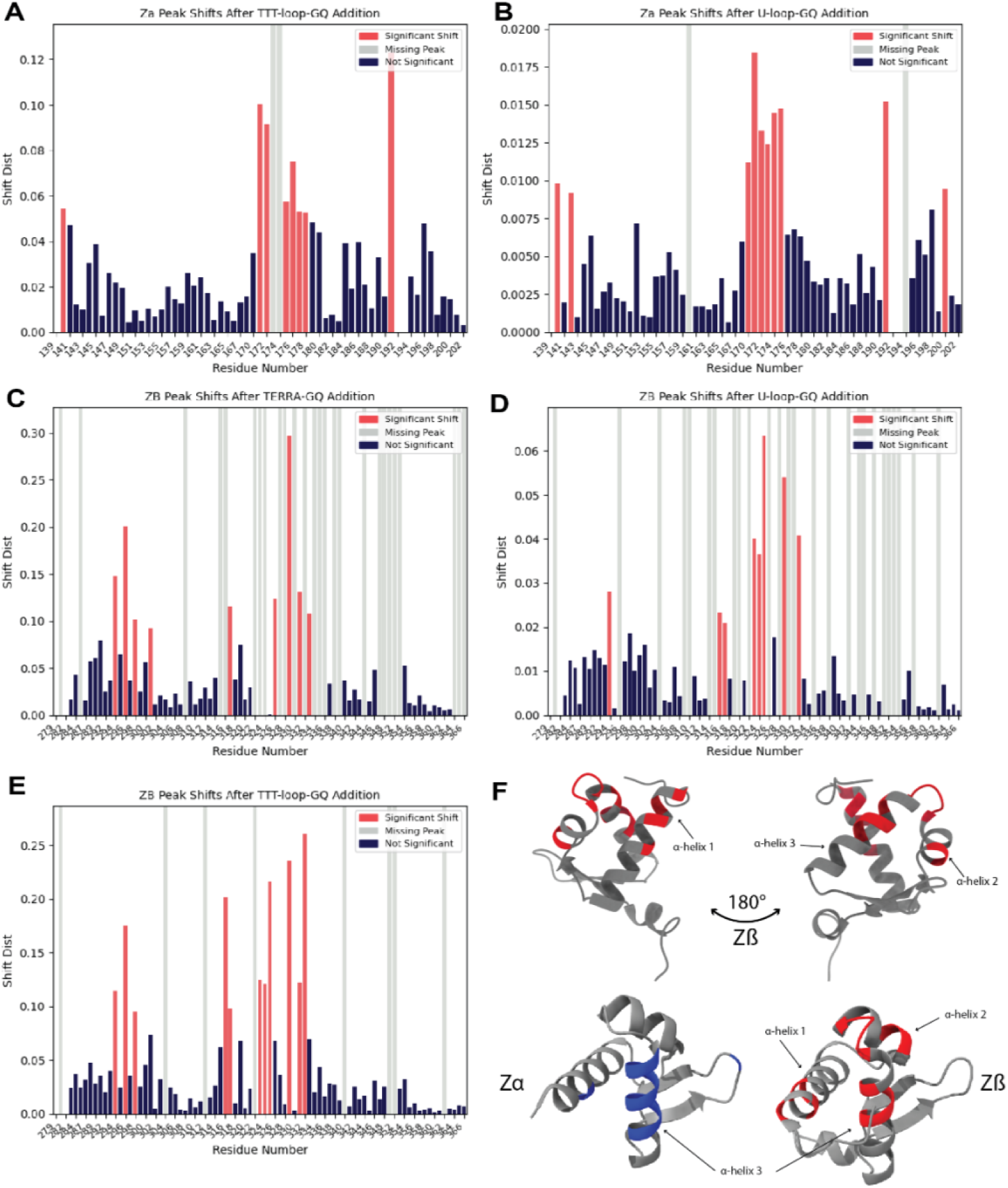
Sites of interaction of ADAR1 Zα and Zβ upon G-quadruplex binding. CSPs obtained from HSQC spectra of Zα upon addition of A) TTT-loop-GQ_DNA_ and B) U-loop-GQ_RNA_ in KCl. CSPs obtained from HSQC spectra of Zβ upon addition of C) TERRA-GQ_RNA_, D) U-loop-GQ_RNA_ in KCl and E) TTT-loop-GQ_DNA_. Red, blue and grey bars represent significant CSPs, non-significant CSPs, and grey bars missing/disappeared peaks. F) Structure of Zβ with residues significantly affected upon addition of GQs highlighted in red (top), and comparison of aligned Zα and Zβ, with significant residues highlighted in blue and red respectively (bottom).

CSP analysis of Zβ interactions with TERRA-GQ_RNA_, U-loop-GQ_RNA_, and TTT-loop-GQ_DNA_ indicates primary affected residues across these three constructs being: Leu294, Met296, Glu298, Leu316, Asn317, Gly323, Leu324, Thr325, Asp329, Asn331, and Ala332, which are affected in binding to at least two of the three GQs. Residues Glu301, Lys326, and Val333 show significant shifts only in TERRA-GQ_RNA_. These residues do shift in both other constructs, but not to a significant level Finally, Arg328, which was hypothesized to play a significant role in GQ interaction, disappeared completely in all GQ binding spectra, but exhibited large shift distances in both mutant sequences. Taken together, these data indicate that the Zβ G-quadruplex binding interface is formed primarily by α-helices 1 and 3, with contributions from portions of α-helix 2 and the intervening loop between α-helices 2 and 3.

Intriguingly, a comparison of the Z-NA binding interface of Zα to the same region of Zβ implicates several critical residues in Zα that diverge in Zβ resulting in an inability to bind Z-NA, two of which being Arg174/Ala332, and Tyr177/Ile335^20^. The most critical residue implicated in Zβ’s lack of Z-NA binding is Tyr177/Ile335; Ile335 peaks have disappeared in Zβ binding to several GQs (Fig. S5B), and present slight changes in other GQs and the mutant sequences. Ala332 is also one of the most strongly shifting residues across all constructs, suggesting that it is critical for recognition of both GQs and single-stranded RNAs. Despite that Zβ shares a common binding interface for both NA folds, α-helix 1 may contribute additional GQ specificity, as this region is more perturbed by GQs than by mutants (Figs. 7C–E, S7F, S9B). Overall, both ZBDs utilize similar regions within α-helix 3, but Zβ additionally incorporates α-helices 1 and 2 into its G-quadruplex binding surface (Fig. 7F).

## 5 Discussion

While it is well established that Zα domains of ADAR1 and ZBP1 bind nucleic acids that are prone to form Z-DNA and Z-RNA (Fig. 8A), it has also been shown by NMR that the Zα domain of ADAR1 binds an oncogenic c-Myc DNA G-quadruplex^37^. However, no binding partner of the structurally homologous Zβ domain of ADAR1 has been identified to date. Based on AlphaFold^51^ and molecular dynamics simulations, we recently suggested that the Zβ domain of ADAR1 targets substrates by recognizing GQs^49^. Here, we significantly extended the range of experimentally confirmed Zα domain/G-quadruplex interactions by demonstrating that the Zα domains of both ADAR1 and ZBP1 bind to both DNA and RNA GQs. We provided the first direct, in vitro experimental evidence for Zβ domain binding to select G-quadruplex RNAs and DNAs, as well as for interactions between ZBDs and ssRNA. We categorized interactions in Table 4. In general, each ZBD interacted with all NA constructs tested in this study. However, the interaction fingerprints differ across constructs; our binding affinity studies show that peak intensity changes correlate with tighter binding, whereas peak shifts indicated weaker affinity. The results suggest that all ZBDs specifically bind to GQs.

**Figure 8:**
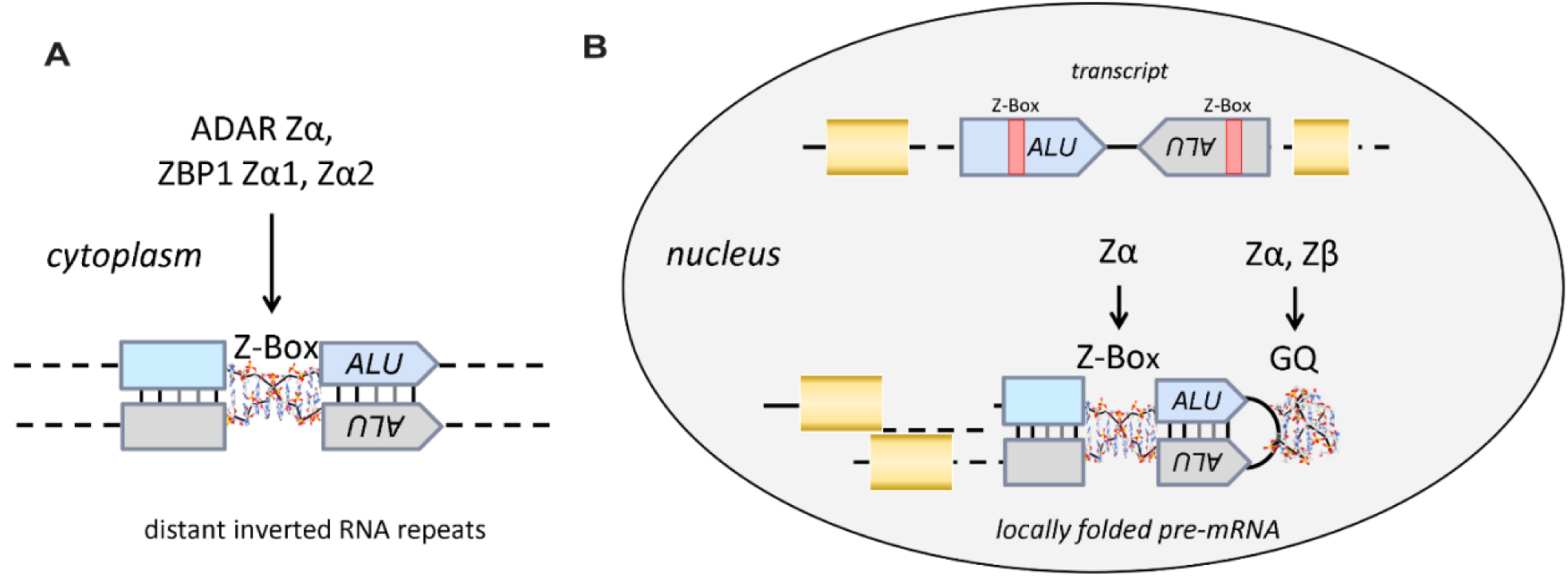
The physiological ligands of ZBPs involved in host defense against retroviral elements. A) Zα domains of ADAR1 and ZBP1 can engage Z-prone regions in dsRNAs found in the cytoplasm, including the Z-Box (a Z-NA prone region) in inverted-repeat ALU elements (IR-ALU). B) Proposed model for ZBD engagement of alternative nucleic acid folds in the nucleus. The Zβ domain binding to GQs in nascent transcripts could potentially localize ADAR1 to edit these RNAs and modulate splicing. GQs may also form in the loop separating the two arms of the IR-ALU. ADAR1’s Zα domain and ZBP1’s Zα1 and Zα2 domains can also bind GQs, but with lower specificity and through a different binding mode compared to Zβ.

We demonstrated that ZBDs interact with both parallel and anti-parallel structures. The antiparallel TTT-loop-GQ_DNA_ exhibited extensive peak shifting, with a notable lack of peak disappearance in contrast to our other GQs. Interactions unique to the TTT-loop-GQ, such as certain Zα peaks shifting in different directions, could potentially be attributable to it being the only DNA GQ and the only anti-parallel GQ examined in this study. Additionally, Zα exhibited a preference for ssRNA over the TTT-loop-GQ DNA, whereas Zβ had a preference for the GQ over the ssRNA. These results indicate that ZBDs recognize both DNA/RNA and parallel/antiparallel GQs through different binding modes.

Additionally, we examined the binding interface used by ADAR1 ZBDs to recognize GQ. Overall, our data agree with the previously proposed regions of interaction²⁷. Zα engages GQ using essentially the same surface that it uses to bind Z-NA, namely α-helix 3 and the β-wing. In contrast, Zβ interacts with GQs primarily through α-helices 1 and 3 and the loop between α-helices 2 and 3 (Fig. 7F). These data provide preliminary evidence that residues within the α-helix 1 region of Zβ contribute to preferential GQ recognition, as Zα does not utilize α-helix 1 in its binding surface. Furthermore, Zβ consistently shows significant shifts in non-polar, aliphatic residues such as isoleucine and alanine; these residues are not typically involved in nucleic acid-binding interactions with B-DNA or A-RNA. However, our observations are consistent with those for the yeast RAP1 transcription factor, which has been crystallized with both B-DNA containing its cognate binding site and a parallel-strand GQ^63,64^. The same RAP1 helix interacts with both conformations, but through different faces. The charged face binds to B-DNA and the hydrophobic face to the hydrophobic end caps of the GQ in a similar manner by which small molecules have been shown to bind GQs^41^. Likewise, we propose that non-polar residues in the ZBD α-helices are hydrophobically interacting with the top or bottom face of GQs, with specific charged residues (like Arg328) contributing to the electrostatic interactions.

Two Zβ residues likely involved in binding to GQs, Ala332 and Ile335, are the positional equivalents of Arg174 and Tyr177 in Zα. Loss of these specific side chains explains Zβ’s inability to bind Z-NA. Their strong involvement in GQ binding suggests that evolution of the Zβ domain may have favored GQ specificity at the expense of Z-NA recognition. The precise functional role of these residues remains unknown, and further structural work is needed to characterize the binding interface better.

The findings support a model we previously proposed^49^ (Fig. 8B), in which the Zβ domain binds GQ formed co-transcriptionally by ALU repeats and other repeats with a GQ motif. The model further predicts that these GQs promote the localization of ADAR1p110 and the editing of dsRNAs formed by ALU inverted repeats, which might otherwise result in detrimental outcomes due to mis-splicing of the pre-mRNA. The recognition of GQs by ADAR1p110 may also promote the resolution of telomeric R-loops^38^. Of note, similar to A-to-I edits, GQs are enriched in human RNA intron splice sites^65^. The role of GQ recognition by ADAR1 in these processes may extend beyond dsRNA editing, as catalytically dead ADAR1 can still alter splicing patterns in mice^53^. Similar to Z-NA recognition by Zα, this mechanism may defend the host against retroelements and exploits the structures they form for other purposes. In the case of GQ, Zβ potentially helps regulate alternative splicing, miRNA processing, and exon recoding. An assessment of A-to-I edits suggested that the associated GQs were usually less than 175 bases from the editing site^49^. This distance is much less than the 7 kb separations possible for ALU inverted repeats that pair to form editing substrates^28^.

Our results also provide new insights into the various roles played by the Zα domain. We present the first experimental evidence for direct binding of a Zα domain to TERRA, extending beyond the previously reported co-localization of ZBP1 to TERRA^34^. Indeed, both the Zα and Zβ domains of ADAR1 show a preference for the TERRA-GQ_RNA_. To confirm specificity for GQ, we examined Zα and Zβ binding to the TERRA-mut_RNA_, in which tetrad formation was disrupted by replacing guanine with cytosine. The affinity for these substrates was an order of magnitude lower than for the GQ fold.

Our findings highlight the distinct modes by which Zα and Zβ target ADAR1 to different sites in the cell, enabling a multi-layered host defense against retroelements and pathogens. GQs can originate from a ssRNA produced during active transcription to localize ADAR1 to inverted-repeat ALUs that are then edited. Alternatively, ssRNAs that form GQs may localize ADAR1 to modulate splicing in an editing-independent manner^53^. These events affect the processing of pre-mRNAs and occur in the nucleus. In the cytoplasm, the formation of Z-RNAs is promoted by helicases and by torsional strains that can be found in stress granules. ADAR1p150 has been shown to localize to stress granules via its Zα domain to negatively regulate immune responses against self-RNAs^11–13,66–68^. As ADAR1p150 also undergoes nucleocytoplasmic shuffling, GQ-forming sequences can be targeted by this isoform in the nucleus. In this scenario, GQ binding by either Zα or Zβ localizes ADAR1 to actively transcribing genes, allowing capture of nascently transcribed RNAs and providing time for folding of ADAR dsRNA intron/exon editing substrates^49^.

ZBP1 Zα1 and Zα2 also engage short GQs (U-loop-GQ_RNA_ and ALU-GQ_RNA_). In general, Zβ was consistently more responsive to variation in GQ structure than other ZBDs. These findings are notable, as they indicate that while both ADAR1 ZBDs can bind GQs, Zβ displays greater apparent structural specificity for the GQs studied. This specificity, along with the replacement Arg174 and Tyr177 of Zα by Ala332 and Ile335 in Zβ, suggests that this domain is better optimized for the parallel-stranded GQs formed by RNAs. The amino-acid substitutions likely facilitate binding to the hydrophobic GQ endcaps. Zα domains also recognize GQ folds. It is likely that binding of Zα1 and Zα2 to GQ positions ZBP1 to capture viral Z-RNAs as they fold. Otherwise, production of high-affinity GQ RNA ligands for Zα1 and Zα2 would enable viruses to inhibit ZBP1-dependent activation of cell-death pathways.

As a final layer of defense, the localization of ZBDs to stress granules is consistent with the phase separates we observe. Studies on liquid-liquid phase separation induced by Zα domains help drive the conversion of A-RNA to Z-RNA and potentially drive anti-viral responses^70,71^. Although this study provides evidence for interactions between the ZBDs of both ADAR1 and ZBP1 with GQs, as well as GQ-specific binding by Zβ, the significant precipitation observed imposes limitations on our results. Several questions regarding the nature of these interactions remain. Specifically, it remains challenging to disentangle the effects of precipitation/phase separation from the actual binding interactions. Several constructs result in the near-complete disappearance of resonances in NMR spectra, precluding quantitative assessment of binding affinity or conformational dynamics and thereby limiting interpretation to a binary presence/absence of binding. Furthermore, NMR requires working at protein and nucleic acid concentrations much higher than those encountered in vivo, raising questions about how our findings translate to an in vivo setting. Nonetheless, this work provides compelling evidence of interactions between ZBDs and GQs and, more intriguingly, a potential answer to the long-standing question about the functional roles of the highly conserved ADAR Zβ domain.

## Funding

This project was supported by NSF grant #2153787 to B.V., and NIH grants R01 GM150642 to B.V., R35GM156171 to B.V., University of Colorado Cancer Center Support Grant P30 CA046934, and Biomedical Research Support Shared Grant S10 OD025020-01.

## Notes

### Competing Interest Statement

The authors have declared no competing interest.

