## Supplemental Information for "Zα and Zβ domains of ADAR1 and ZBP1 bind to G-quadruplexes with low micromolar affinity"

The following information is supplemental and cannot be contained in the main body of the paper.

**1 Analysis of nucleic acid structure**

GQ nucleic acids exhibit characteristic CD spectra, wherein both parallel DNA and RNA GQ have a positive peak at 260 nm, and a negative peak at 240 nm. Antiparallel GQ (usually only formed by DNA) have positive peaks at 295 nm and 240 nm, and a negative peak at 260 nm[1]. These spectra differ from A-form RNA and ssRNA (maximum and minimum at 265-270/210-220 nm), B-form DNA and ssDNA (maximum/minimum at 275-280/245-250 nm), and Z-DNA and Z-RNA (295/260 nm)[2]. These values are described in Table S1.

By CD, we verified that each of our expected GQ sequences was folding in solution (Fig. S1A). TERRA-$GQ_{RNA}$ showed the strongest GQ spectral characteristics, with a significant positive peak at 260 nm, and a negative one at 240 nm. As TERRA is a well-known GQ, we used this sequence to confirm that the other expected GQ RNA sequences, ALU-$GQ_{RNA}$ and U-loop-$GQ_{RNA}$, were properly folded. The CD spectrum of ALU-$GQ_{RNA}$ closely matches that of TERRA-$GQ_{RNA}$, with a strong positive peak at 260 nm and a negative at 240 nm, but with a lower maximum ellipticity at 260 nm. The U-loop-$GQ_{RNA}$ showed a similar CD pattern indicative of a folded parallel GQ, but with slightly lower maximum ellipticity than TERRA-$GQ_{RNA}$ or ALU-$GQ_{RNA}$. These results indicate that we have successfully folded each of our RNA GQ sequences in GQ-promoting buffer. This is in contrast to our negative controls, TERRA-$mut_{RNA}$ and ALU-$mut_{RNA}$, as their sequences cannot form GQ structures. As expected, their CD spectra show a positive peak at 275 nm, and do not have a negative peak at 240. This confirms that our $GQ_{RNA}$ sequences are folding into parallel GQ structures while our mutated sequences are not. We also confirmed that the TTT-loop-$GQ_{DNA}$ forms an antiparallel GQ, exhibiting characteristic positive peaks at 295 and 240 nm, and a negative peak at 260 nm (Fig. S1B).

### 1.1 G-quadruplex melting curves

To further characterize the folding of our RNA and DNA constructs into G-quadruplexes, we examined melting temperatures under various buffer conditions. As RNA G-quadruplexes are very stable and known to melt at temperatures upwards of 80 °C[3,4], by altering the predominant ion in solution between $K^+$, $Na^+$, and $Li^+$, we can visualize a difference in melting temperature (Fig. S1C). While lithium disfavors or even prevents its formation, sodium moderately and potassium strongly favor its formation[3,5].

We observed high melting temperatures for the three confirmed GQ-forming RNA constructs in KCl buffer: both ALU- and U-loop $GQ_{RNA}$ had melting temperatures above 90 °C, and TERRA-$GQ_{RNA}$ at 80 °C. These values are significantly higher than those of the non-GQ-forming TERRA-$mut_{RNA}$, which melted at 45 °C. Furthermore, the melting temperatures decreased accordingly with the GQ ion ‘hierarchy’ ($K^+ > Na^+ > Li^+$). ALU-$GQ_{RNA}$ showed melting temperatures of 73 and 64 °C in NaCl/LiCl, while U-$loop_{RNA}$ demonstrated melting temperatures of 78 and 69 °C, respectively. Additionally, TERRA-$GQ_{RNA}$ exhibited a melting temperature of 45 °C in the presence of LiCl. In contrast, TERRA-$mut_{RNA}$ exhibited equivalent melting temperatures of 45 °C between KCl and LiCl buffers (Fig. S1C). These results confirm that each RNA GQ construct forms a G-quadruplex in solution, and that by varying the predominant ion in solution, we can alter the stability of the G-quadruplex structure.

### 1.2NMR analysis of G-quadruplexes

NMR is a powerful tool orthogonal to CD spectroscopy to determine nucleic acid structure. In the G-tetrad, there are four hydrogen bonds between the imino proton on N1, and the neighboring O6 on the neighboring guanine (Fig. S1D, circled red). These hydrogen bonds are stable and prevent hydrogen exchange with the solution, allowing characteristic G-quadruplex peaks to be visible in the 10-12 ppm region in a $^{1}$H NMR spectrum. Other RNA structures in which the imino proton is visible show imino peaks further downfield (for example, hairpin RNA has imino peaks in the 13-14 ppm range)[3]. In the TERRA-mut$_{RNA}$ sequence, guanines have been swapped to cytosine, blocking the ability for G-tetrads to form and subsequently allowing N1 hydrogens to exchange with solution readily. Thus, there should be no visible peaks in the 10-12 ppm region for non-GQ RNA. Our 1D NMR analysis of the TERRA-GQ$_{RNA}$ showed strong peaks in the 10-12 ppm range, further confirming its ability to form a GQ structure, while the TERRA-mut$_{RNA}$ had no visible peaks in this range (Fig. S1D).

We also analyzed ALU- and U-loop-GQ$_{RNA}$ via $^{1}$H NMR. Both of these constructs exhibited peaks in the GQ imino region, but with slightly broader peaks than seen with TERRA-GQ$_{RNA}$. This could be a result of a less stable GQ structure, but still indicates the presence of a G-quadruplex. Subsequently, we also examined the effects of altering the predominant buffer ion via NMR. In accordance with our melting results, we observe that for both ALU- and U-loop-GQ, the buffer condition reduces the stability/degree of folded GQ. We see that in high $Na^{+}$ buffer, the imino peaks shift slightly downfield and begin to experience some peak broadening. Furthermore, in $Li^{+}$ buffer, the peaks remain at the same position as $Na^{+}$, but are even broader (Fig. S1). This suggests that not only do we decrease the stability of GQ, but we also shift the equilibrium in solution towards a more unfolded state. We also examined the temperature stability of ALU-GQ$_{RNA}$ in various buffers. In $K^{+}$ buffer, there are no visible changes to the imino peaks between 35-75°C. In contrast, in $Na^{+}$ buffer, we begin to see peak intensity reductions as temperature increases, and the complete disappearance of imino peaks in $Li^{+}$ (Fig. S2).

Taking the CD, temperature melting and NMR data together, we confirmed that all our constructs of interest form G-quadruplexes under our experimental *in vitro* conditions. Placing G-quadruplexes in either KCl, NaCl, or LiCl buffer exhibits variable stability depending on the predominant ion in the buffer, with $K^{+}$ having the strongest GQ-promoting effects, and $Li^{+}$ the weakest. Thus, we can utilize these modulations to characterize GQ stability and to study if Zα and Zβ domains exhibit differential binding to the same RNA sequence depending on its secondary structure.

Table S1: Comparison of characteristic maximum and minimum CD ellipticity values for different DNA and RNA secondary structures.

| | **Positive peak (nm)** | **Negative peak (nm)** | **Construct** |
|---|---|---|---|
| A-RNA, ssRNA | 265-270 | 210-220 | TERRA-$mut_{RNA}$, ALU-$mut_{RNA}$ |
| B-DNA, ssDNA | 275-280 | 245-250 | - |
| Z-DNA, Z-RNA | 295 | 260 | - |
| Antiparallel G-quadruplex | 295,240 | 260 | TTT-loop-$GQ_{DNA}$ |
| Parallel G-quadruplex | 260 | 240 | TERRA-$GQ_{RNA}$, ALU-$GQ_{RNA}$, U-loop-$GQ_{RNA}$ |

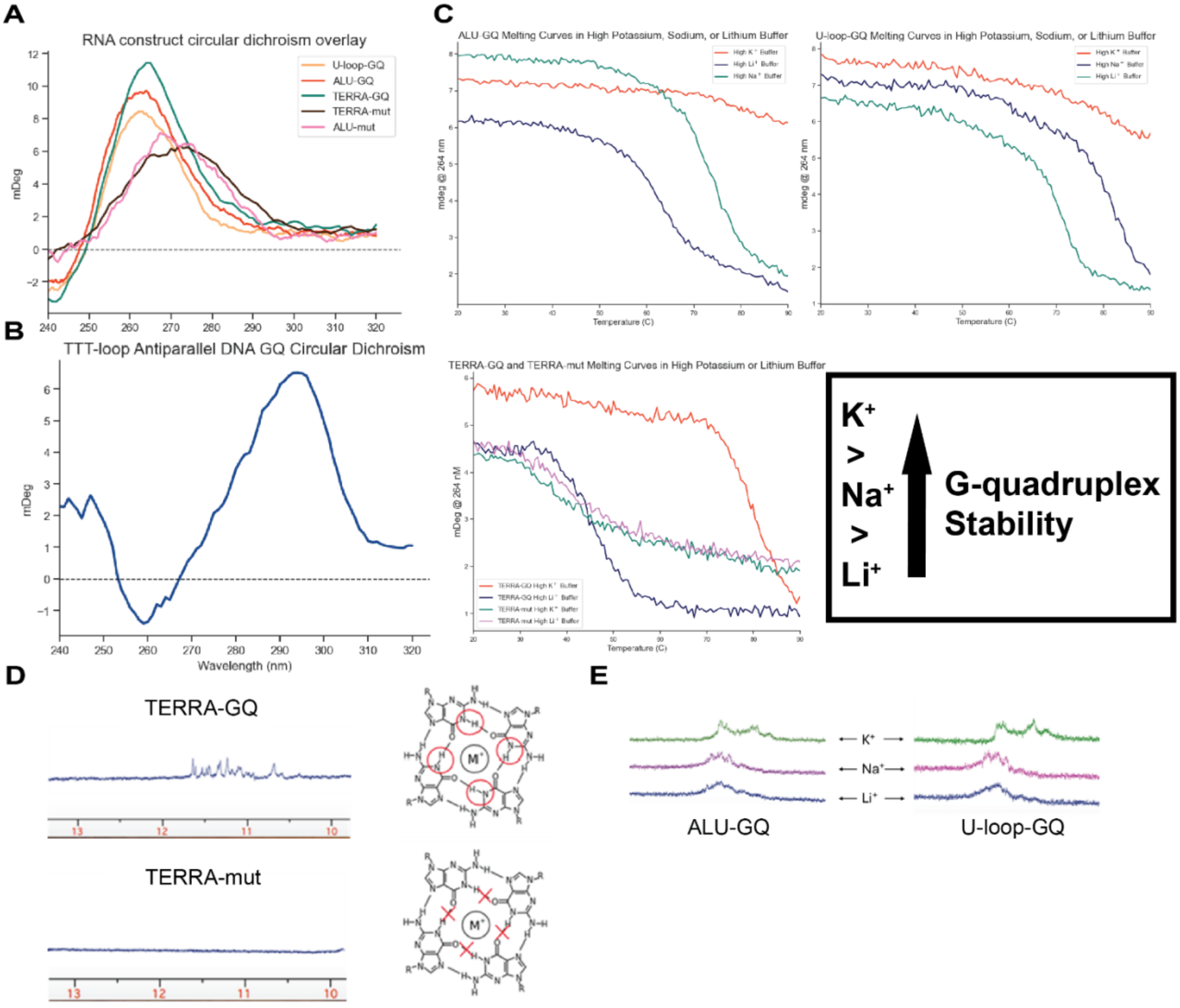


Figure S1: **Assessment of G-quadruplex formation of RNA and DNA constructs**. A) Overlaid CD spectra of RNA constructs, all constructs except TERRA-mut$_{RNA}$ and ALU-mut$_{RNA}$ adopt a parallel GQ structure. B) CD spectrum of TTT-loop-GQ$_{DNA}$ shows peaks typical of antiparallel GQ. C) CD melting curves of ALU-GQ$_{RNA}$ and U-loop-GQ$_{RNA}$ in KCl, NaCl, or LiCl buffer, and TERRA-GQ$_{RNA}$ and TERRA-mut$_{RNA}$ in KCl or LiCl buffer at 35 °C. D) Imino region $^{1}$H NMR shows GQ peaks for TERRA-GQ$_{RNA}$ but no peaks for TERRA-mut$_{RNA}$. E) Comparison of $^{1}$H NMR imino peaks for ALU-GQ$_{RNA}$ and U-loop-GQ$_{RNA}$ in KCl, NaCl, or LiCl buffer at 35 °C.

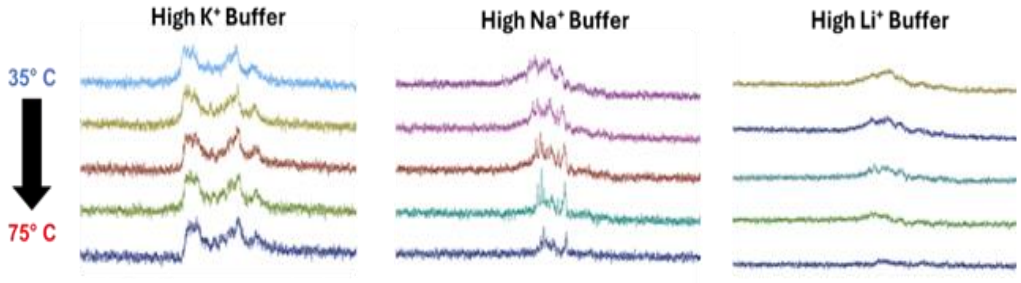


Figure S2: **G-quadruplex melting by NMR.** U-loop-GQ$_{RNA}$ in KCL, NaCl, or LiCl buffer at 35, 45, 55, 65, and 75 °C. Temperature has no visible effect in the $K^+$ buffer, while $Na^+$ exhibits mild temperature sensitivity, as evidenced by peak intensity reduction; the GQ signal is completely abrogated at 75°C in the $Li^+$ buffer.

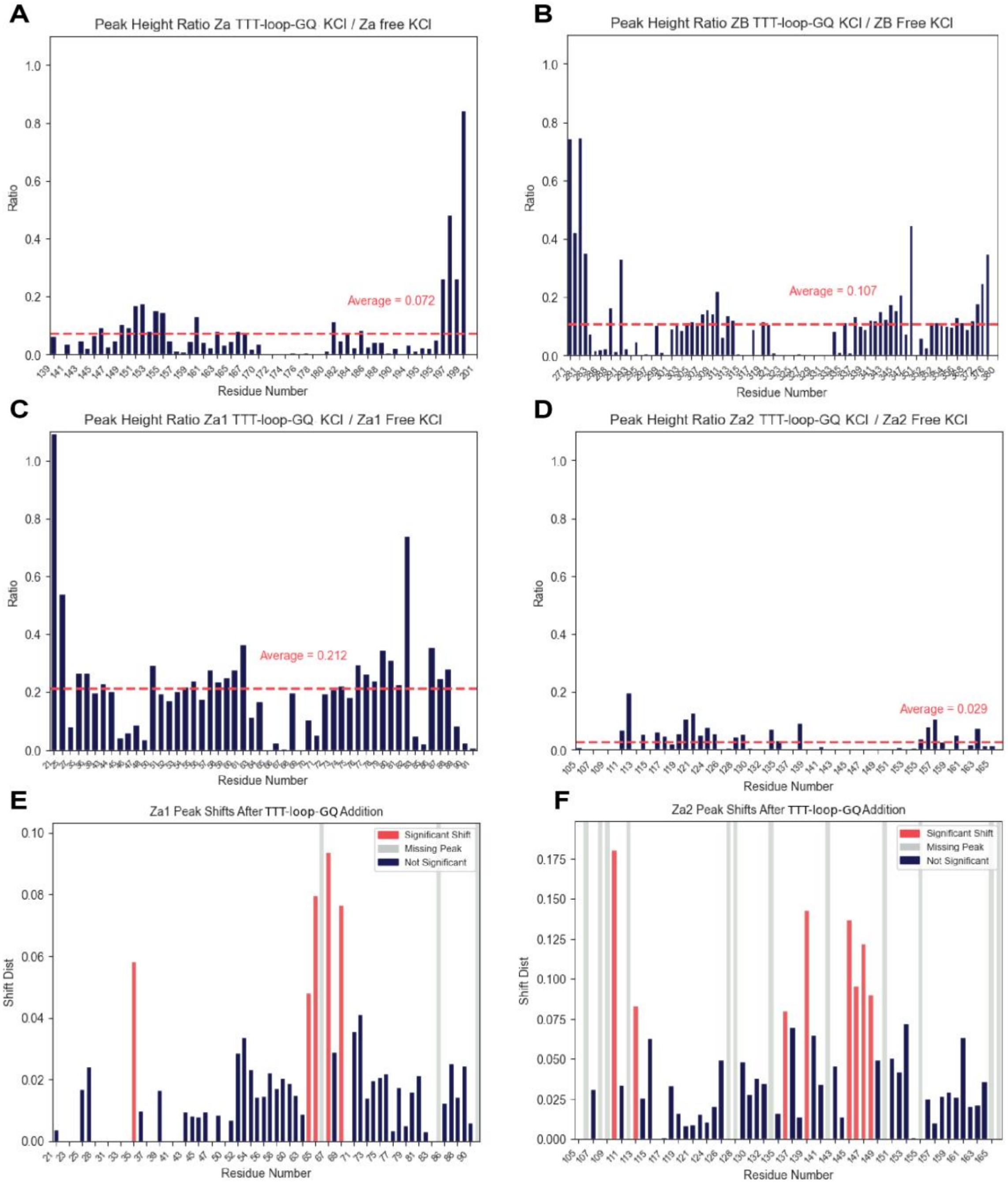


Figure S3: **HSQC peak height ratio plots of ADAR1 and ZBP1 ZBDs binding to TTT-loop-GQ$_{DNA}$, CSP plots of binding with Zα1 and Zα2.** Plots are represented as a ratio of bound (nucleic acid + protein) to free protein. The red line indicates the mean. A-D) ADAR1 Zα, Zβ, ZBP1 Zα1, Zα2 + TTT-loop-GQ$_{DNA}$, respectively. E, F) CSP plots binding with Zα1 and Zα2 respectively.

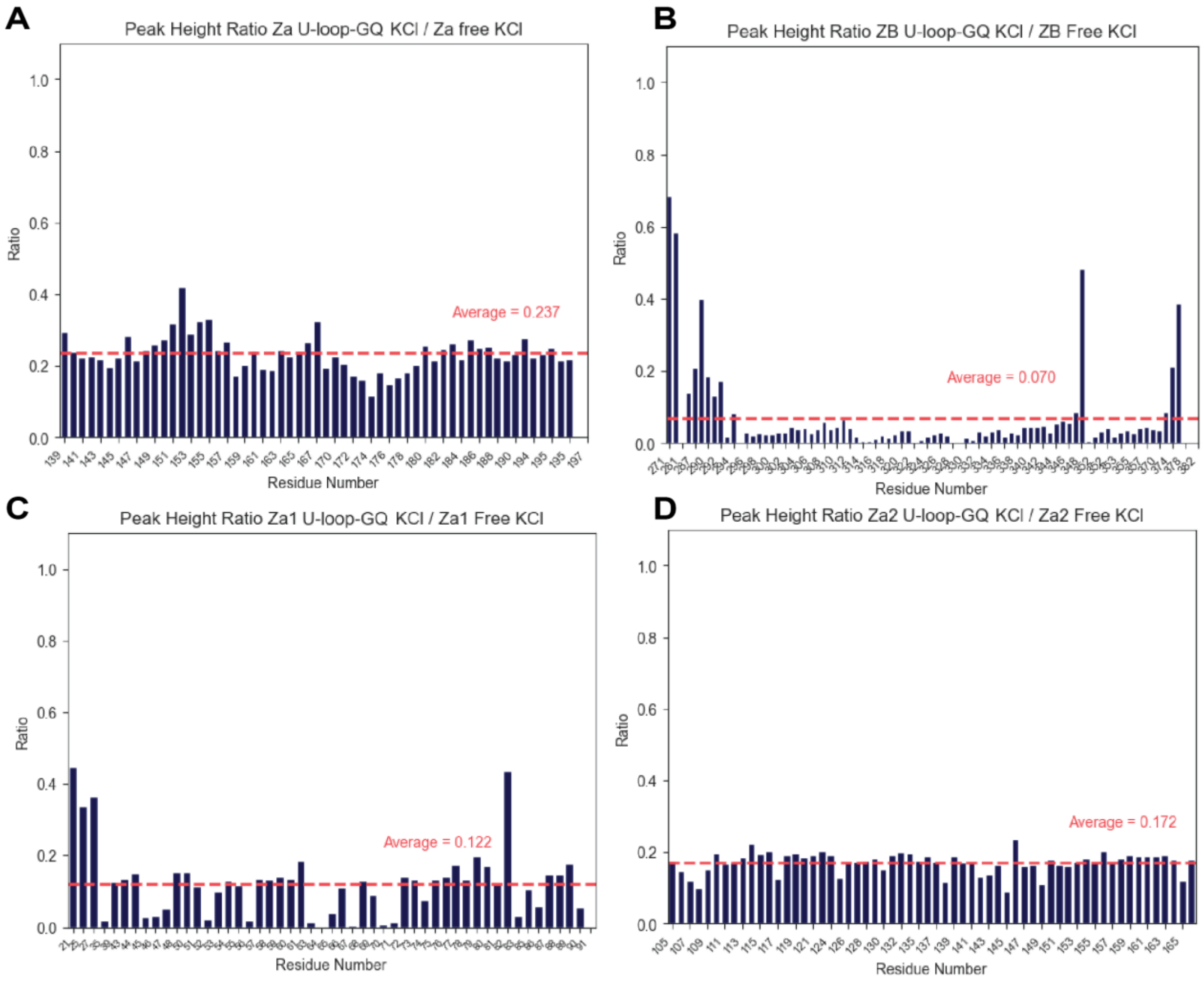


Figure S4: **HSQC peak height ratio plots of ADAR1 and ZBP1 ZBDs binding to U-loop-GQ$_{RNA}$.** Plots are represented as a ratio of bound (nucleic acid + protein) to free protein. The red line indicates the mean. A-D) ADAR1 Zα, Zβ, ZBP1 Zα1, Zα2 + U-loop-GQ$_{RNA}$, respectively.

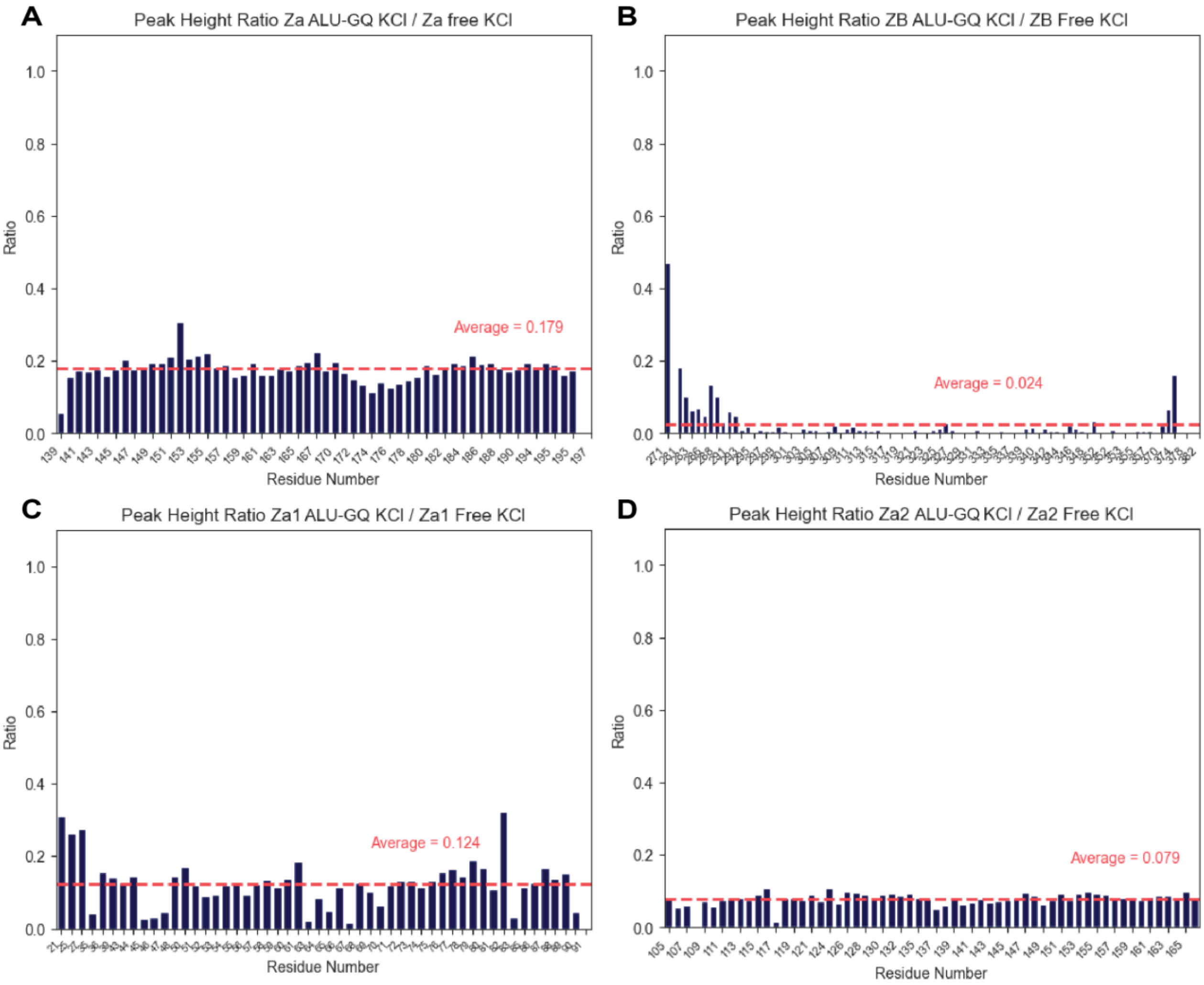


Figure S5: **HSQC peak height ratio plots of ADAR1 and ZBP1 ZBDs binding to ALU-GQ$_{RNA}$.** Plots represented as a ratio of bound (nucleic acid + protein)/free protein. The red line indicates the mean. A-D) ADAR1 Zα, Zβ, ZBP1 Zα1, Zα2 + ALU-GQ$_{RNA}$ respectively.

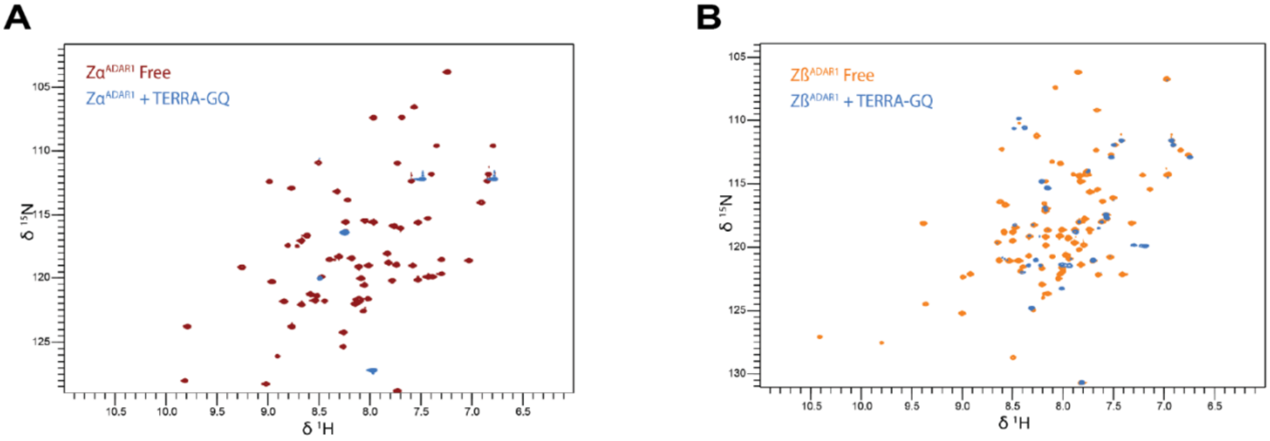


Figure S6: **Comparison of interaction of ADAR1 Zα and Zβ subunits with TERRA-GQ$_{\mathbf{RNA}}$.** Overlaid 2D HSQC spectra of free or bound protein of A) free Zα ADAR1 free and in the presence of TERRA-GQ$_{RNA}$, B) free Zβ ADAR1 free and in the presence of TERRA-GQ$_{RNA}$.

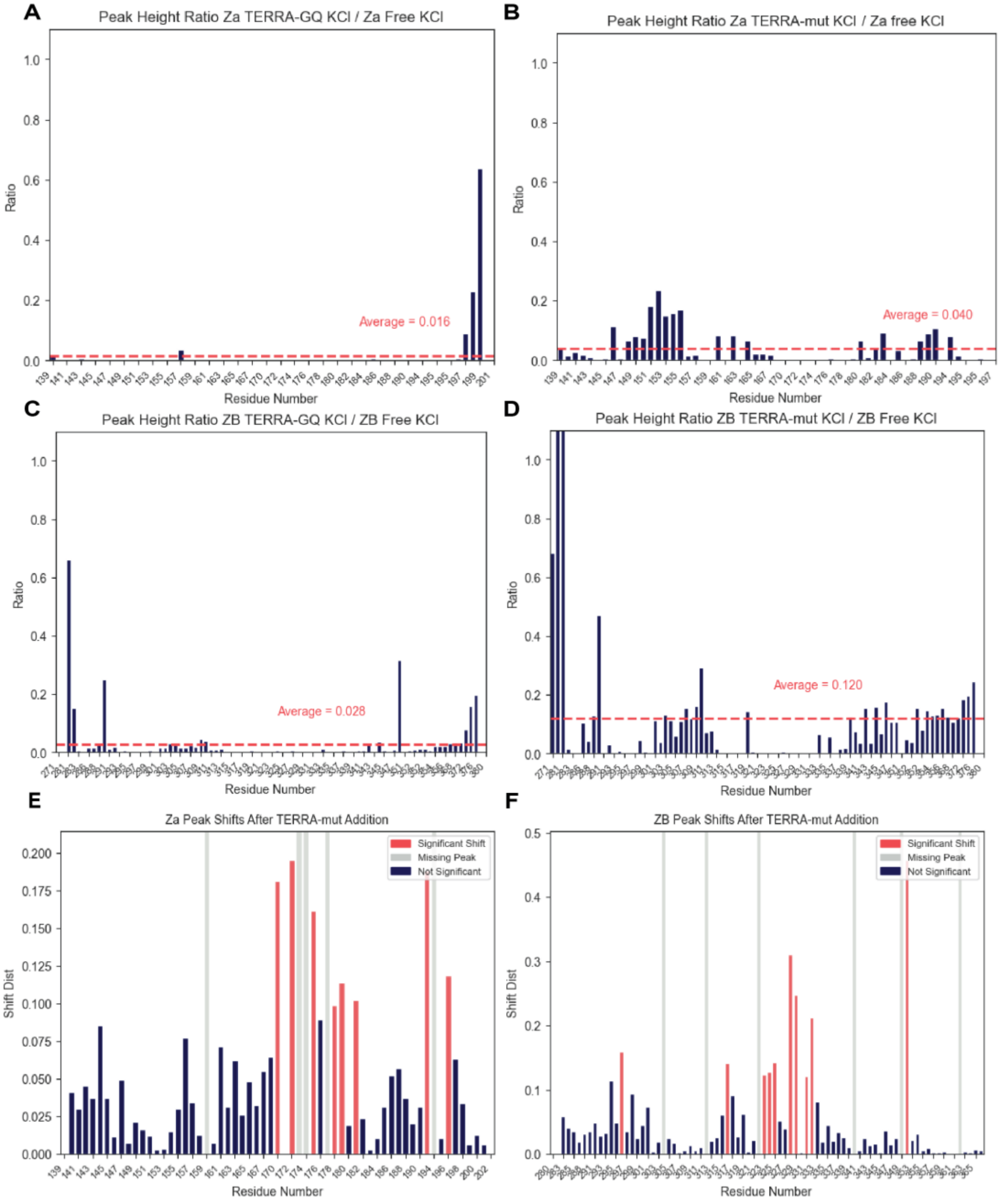


Figure S7: **HSQC peak height ratio plots and CSP plots of ADAR1 Zα and Zβ binding to TERRA-GQ$_{\text{RNA}}$ and TERRA-mut$_{\text{RNA}}$.** Plots are represented as a ratio of bound (nucleic acid + protein)/free protein. A, B) Peak height ratios of Zα + TERRA-GQ$_{\text{RNA}}$ or TERRA-mut$_{\text{RNA}}$. C, D) Peak height ratios of Zβ + TERRA-GQ$_{\text{RNA}}$ or TERRA-mut$_{\text{RNA}}$. E, F) CSPs of Zα and Zβ binding to TERRA-mut$_{\text{RNA}}$, respectively.

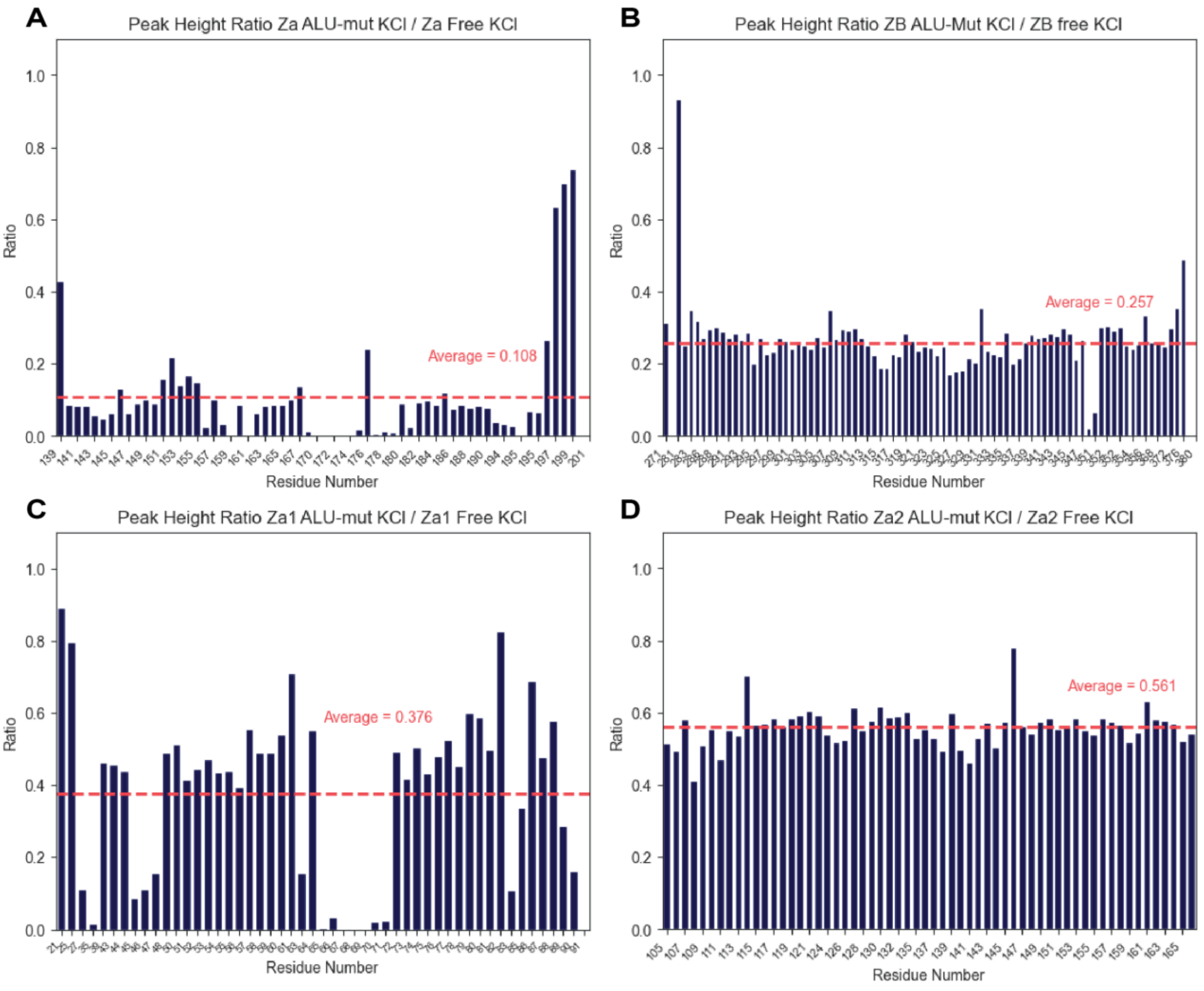


Figure S8: **HSQC peak height ratio plots of ADAR1 and ZBP1 ZBDs binding to ALU-mut$_{RNA}$.** Plots are represented as a ratio of bound (nucleic acid + protein)/free protein. The red line indicates the mean. A-D) ADAR1 Zα, Zβ, ZBP1 Zα1, Zα2 + ALU-mut$_{RNA}$, respectively.

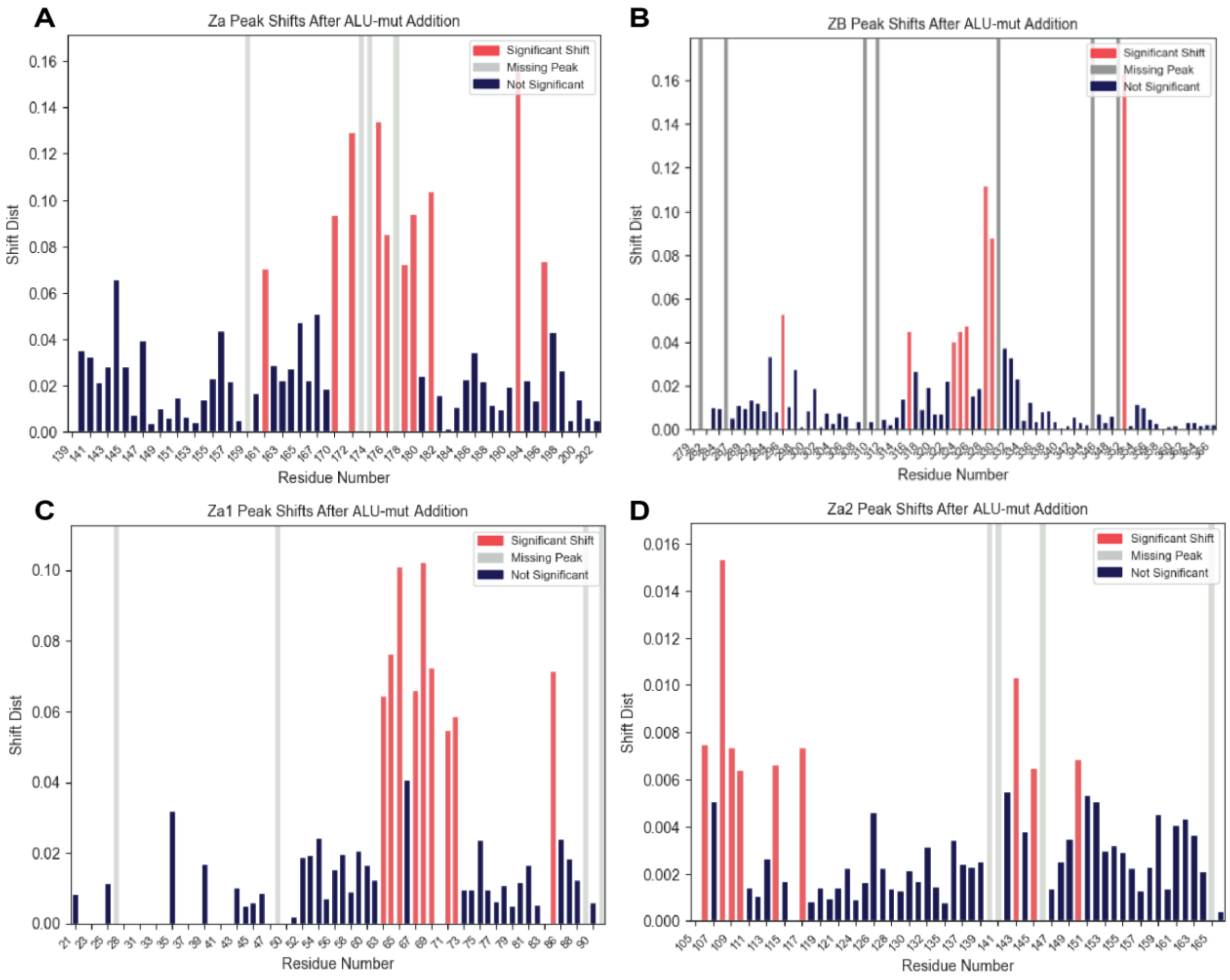


Figure S9: **Chemical shift perturbation plots of ADAR1 and ZBP1 ZBDs binding to Alu-mut$_{RNA}$.** A-D) CSPs of ADAR1 Zα, Zβ, ZBP1 Zα1, and Zα2 binding to Alu-mut$_{RNA}$, respectively.

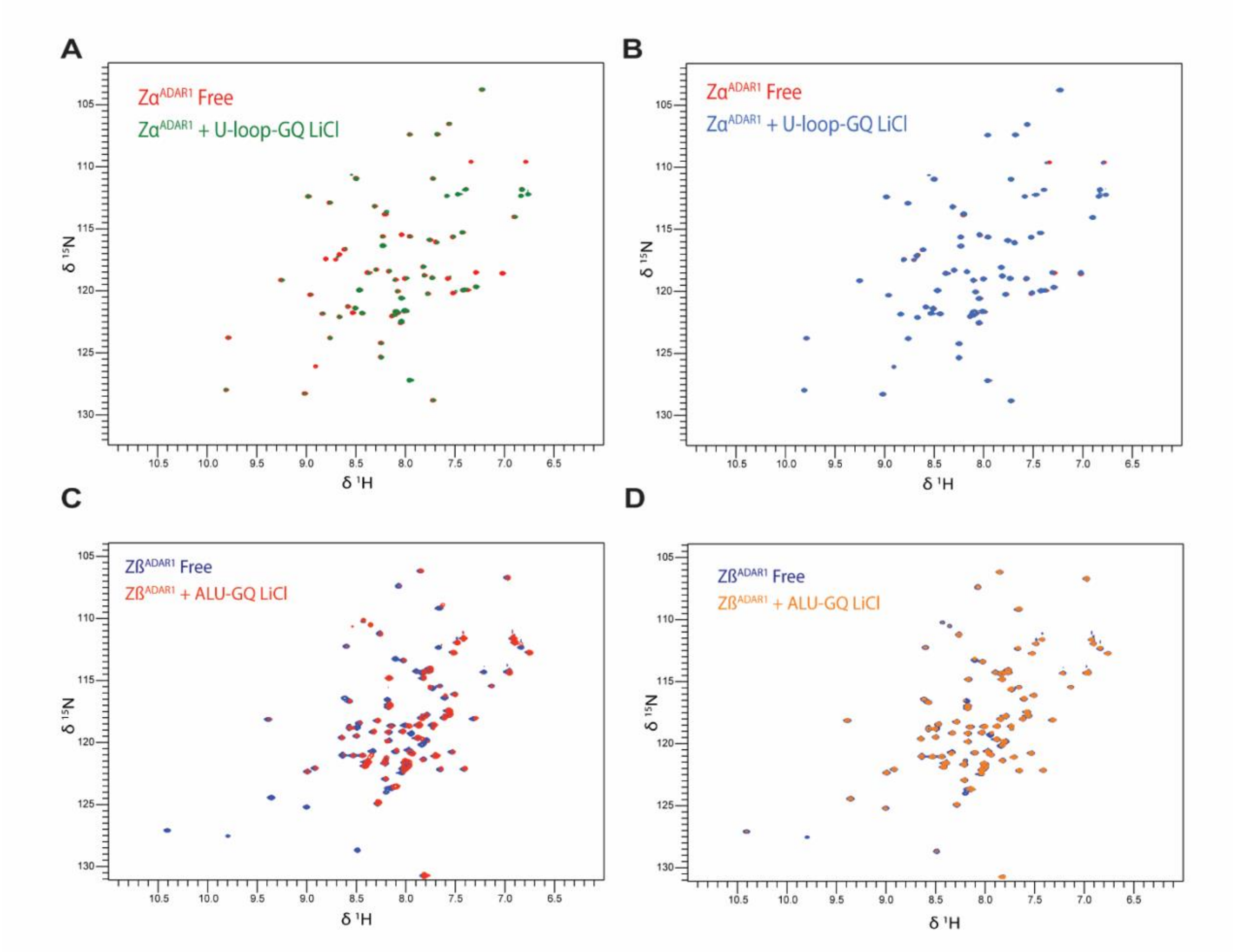


Figure S10: **Comparison of ADAR1 ZBD binding to ALU-GQ$_{RNA}$ in KCl and LiCl buffer.** Overlay of HSQC spectra in high Li$^{+}$ of Zα domain in complex with A) ALU-GQ$_{RNA}$ and B) U-loop-GQ$_{RNA}$. Overlay of HSQC spectra in high Li$^{+}$ of Zβ domain in complex with C) ALU-GQ$_{RNA}$ and D) U-loop-GQ$_{RNA}$.

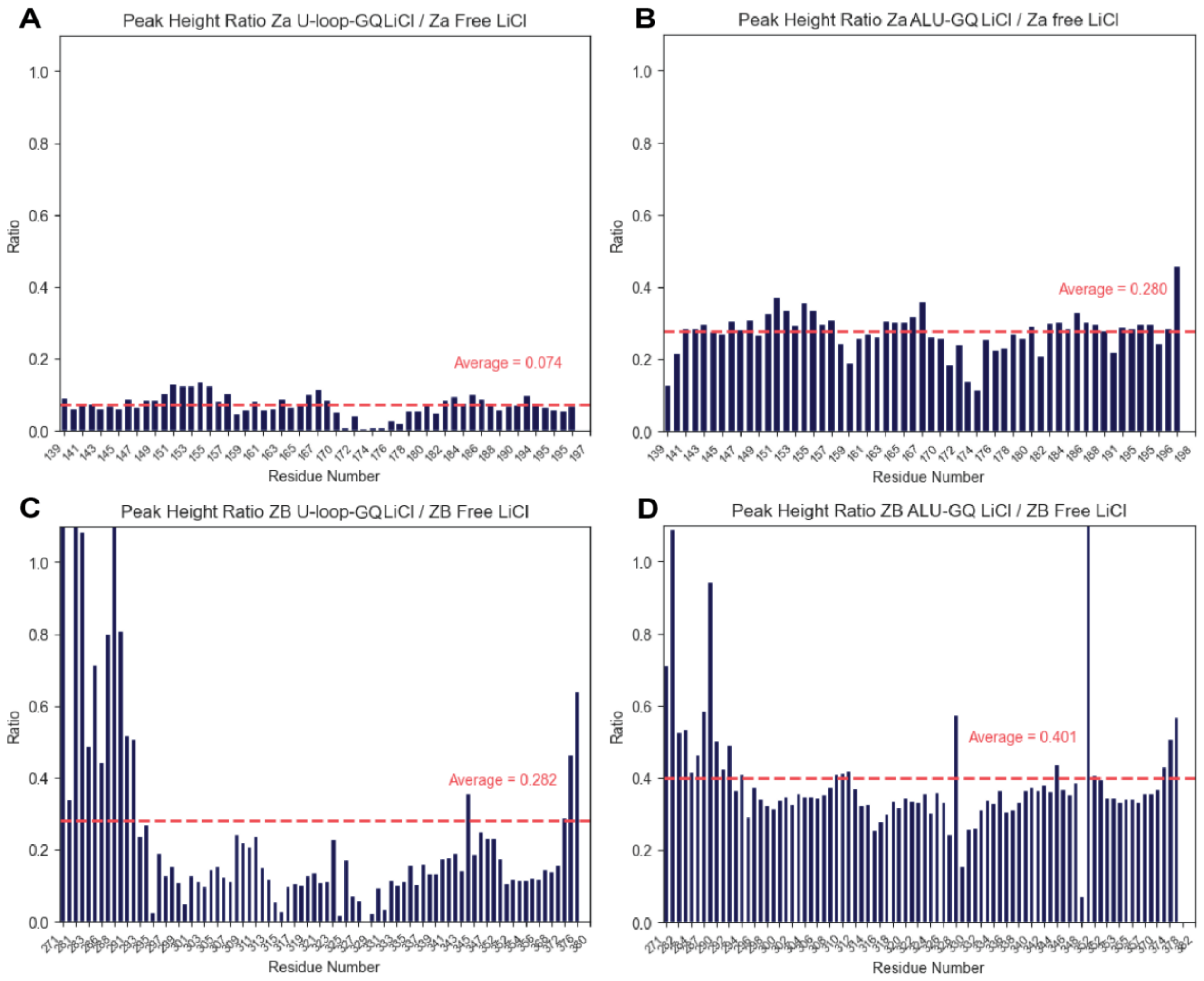


Figure S11: **HSQC peak height ratio plots of ADAR1 Zα and Zβ binding to U-loop-GQ and ALU-GQ$_{RNA}$ in Li$^+$ buffer.** Plots are represented as a ratio of bound (nucleic acid + protein)/free protein. A, B) Zα + U-loop-GQ$_{RNA}$ and ALU-GQ$_{RNA}$, respectively. C, D) Zβ + U-loop-GQ$_{RNA}$ and ALU-GQ$_{RNA}$, respectively.

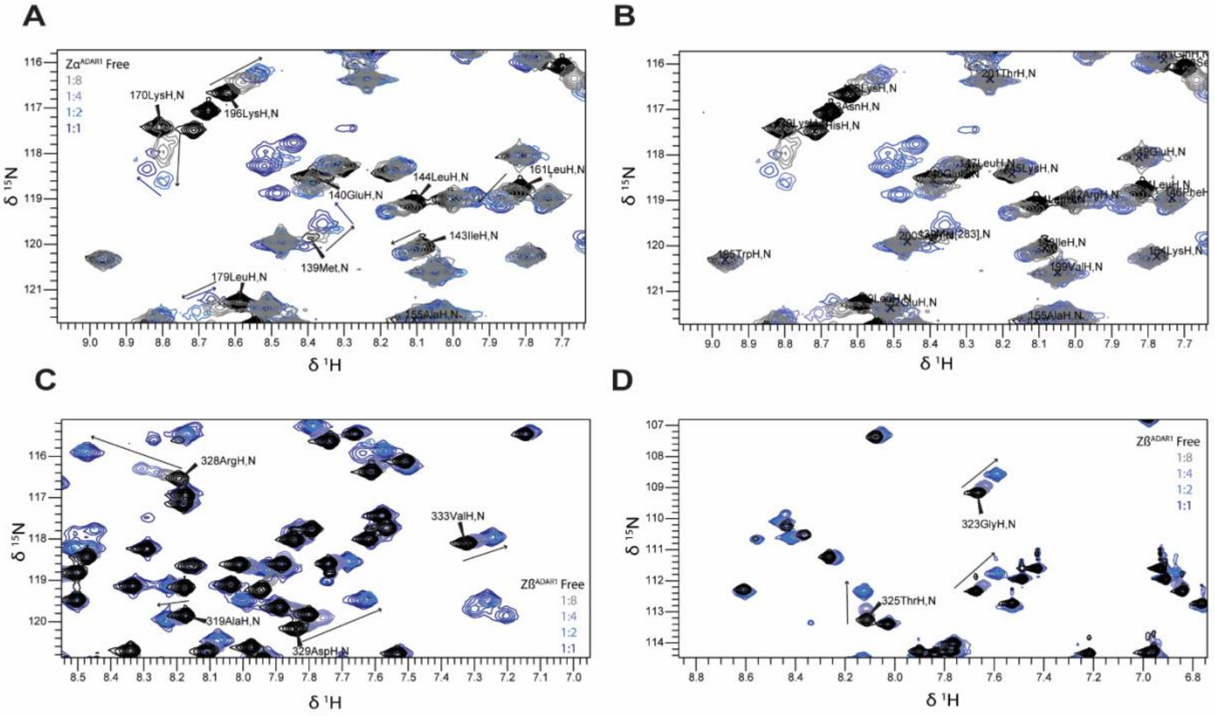


Figure S12: **ADAR1 Zα and Zβ titration series with TERRA-mut$_{RNA}$.** A, B) Extended regions of overlaid HSQC spectra of Zα domain with increasing concentrations of TERRA-mut$_{RNA}$. Concentrations are 200 μM Zα, and 25, 50, 100, and 200 μM TERRA-mut$_{RNA}$. C, D) Extended regions of overlaid HSQC spectra of Zβ domain with increasing concentrations of TERRA-mut$_{RNA}$. Concentrations are 200 μM Zβ, and 25, 50, 100, and 200 μM TERRA-mut$_{RNA}$.

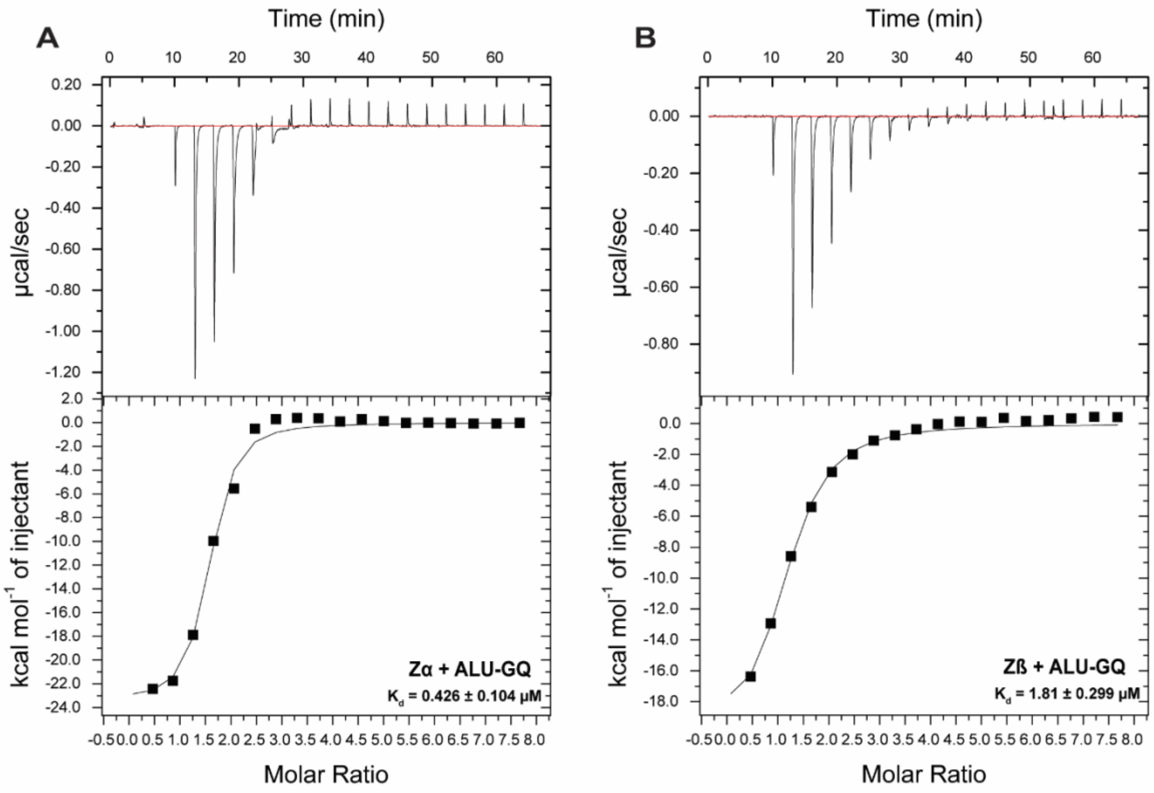


Figure S13: **Full isothermal titration calorimetry spectra of ADAR1 ZBDs binding to ALU-GQ$_{RNA}$.** A) Zα domain + ALU-GQ$_{RNA}$. B) Zβ domain + ALU-GQ$_{RNA}$.
